# Modeling and measuring how codon usage modulates the relationship between burden and yield during protein overexpression in bacteria

**DOI:** 10.1101/2024.11.28.625058

**Authors:** Cameron T. Roots, Alexis M. Hill, Claus O. Wilke, Jeffrey E. Barrick

## Abstract

Excess utilization of translational resources is a critical source of burden on cells engineered to over-express exogenous proteins. To improve protein yields and genetic stability, researchers often use codon optimization strategies that improve translational efficiency by matching an exogenous gene’s codon usage with that of the host organism’s highly expressed genes. Despite empirical data that shows the benefits of codon optimization, little is known quantitatively about the relationship between codon usage bias and the burden imposed by protein overexpression. Here, we develop and experimentally evaluate a stochastic gene expression model that considers the impact of codon usage bias on the availability of ribosomes and different tRNAs in a cell. In agreement with other studies, our model shows that increasing exogenous protein expression decreases production of native cellular proteins in a linear fashion. We also find that the slope of this relationship is modulated by how well the codon usage bias of the exogenous gene and the host’s genes match. Strikingly, we predict that an overoptimization domain exists where further increasing usage of optimal codons worsens yield and burden. We test our model by expressing sfGFP and mCherry2 from constructs that have a wide range of codon optimization levels in *Escherichia coli*. The results agree with our model, including for an mCherry2 gene sequence that appears to lose expression and genetic stability from codon overoptimization. Our findings can be leveraged by researchers to predict and design more optimal cellular systems through the use of more nuanced codon optimization strategies.

## 1 Introduction

Bacterial cells have evolved metabolic pathways and gene regulatory networks that achieve high fitness in their natural environments. When bacteria are genetically engineered, their new functions can impact the availability of cellular resources, induce stress/toxicity, or otherwise interfere with growth and survival [1–4]. The fitness cost of adding an engineered DNA construct to a host cell is referred to as its burden. Constructs with high burden can be evolutionary unstable or even unclonable if cells with mutations that alleviate that burden rapidly arise and take over a population [5–7]. Burden has limited the performance and functional lifetimes of cells with complex genetic circuits and metabolic engineering at both laboratory and commercial scales [6, 8–11].

Engineered cells must distribute gene expression resources between native and added functions. Depletion of RNA polymerase molecules, nucleotides, ribosomes, tRNAs, amino acids, or other transcription and translation factors by genes in an engineered DNA construct will reduce expression of a cell’s genes and slow its replication. Of these factors, ribosomes are most commonly cited as the limiting resource when proteins are overexpressed in engineered *E. coli* cells [2, 12–16]. As an mRNA is produced from an engineered DNA construct, ribosomes that would otherwise be allocated to make a cell’s native proteins needed for replication are instead diverted to produce the exogenous protein, causing gene expression burden.

Because the concentrations of different tRNA species in cells and the rates at which they are charged by aminoacyl-tRNA synthetases vary, codon usage affects the expression of proteins by impacting translational efficiency, the rate at which ribosomes move along an mRNA as they synthesize the encoded protein [17–22]. Codon usage bias in genes can affect the degree to which their production impacts native gene expression and therefore the burden of protein overexpression [2, 23]. It is common for researchers to adopt codon optimization strategies to achieve higher expression levels of an engineered construct by improving its translational efficiency. Common methods include maximizing Fraction of Optimal codons (FOP) [24] or Codon Adaptation Index (CAI) [25] scores. Alternatively, newer global codon harmonization [26] and codon health index (CHI) [22] metrics can be used. The importance of optimizing codon usage to achieve high levels of protein expression is also evident from how deoptimizing codon usage in the major capsid protein of bacteriophage T7 markedly decreases its fitness [27] and from how other bacteriophages encode their own tRNAs to reprogram host codon usage [28]. Similarly, higher yields of recombinant proteins can be achieved in *E. coli* strains engineered to overexpress rare tRNAs [29].

Researchers must have a robust understanding of cellular resources and when they become limiting to engineer cells that efficiently and stably carry out new functions. To improve our understanding of the relationship between codon usage bias and burden from protein overexpression, we simulated and experimentally characterized overexpression of proteins from genes with a range of different codon usage biases in *E. coli*. We found that there is a negative linear relationship between protein production and burden, regardless of the level of codon (de)optimization, and that codon usage biases which deviate from a cell’s charged tRNA availability exacerbate the fitness cost of overexpression.

## 2 Results

### 2.1. Gene expression burden depends on codon usage in simulations

Prior studies have shown that there is an inverse relationship between exogenous protein overexpression and the growth rate of a bacterial cell [2, 12, 30]. Less is known about how this relationship is influenced by codon usage. Expression of an mRNA encoding an exogenous gene that has less optimal codon usage is expected to be more burdensome to a cell because its translation requires more rare tRNAs, which are a limiting resource. To test this hypothesis, we developed a computational model for protein overexpression using Pinetree [31], a stochastic gene expression simulation framework. Pinetree models translation by explicitly tracking ribosomes as they move along mRNAs. We updated Pinetree to also simulate dynamic tRNA recycling, in which charged tRNAs are depleted by translation of their cognate codon and then replenished by aminoacyl-tRNA synthetase reactions. These simulations also take into account how tRNA abundance affects the kinetics of translating ribosomes. The rate of decoding a codon and translocating to the next codon is proportional to the concentration of the relevant charged tRNA species.

In the simulations, we used a mixture of two classes of tRNAs, corresponding to optimal and non-optimal codons. We set up simulations so that the concentration of one of the tRNAs is greater than the other, which makes its codon optimal in terms of codon usage bias. Specifically, we set the relative proportions of these two tRNAs to 70% and 30% of the total tRNA pool, respectively. Growth of cells is proportional to the production of a generic stand-in for all cellular protein (CP), which is encoded by an mRNA that has codon usage biased toward the optimal codon. Then, we add an mRNA encoding an exogenous protein with its own codon usage bias. We simulated different levels of overexpression of the exogenous protein by varying the rate of translation initiation on its mRNA. For the overexpressed protein (OEP), we simulated five different mRNA constructs, with codon usage biases ranging from 10% to 90% optimal. The fitness burden the simulated cell experiences from OEP production is equal to how much its presence decreases expression of CP relative to its baseline level when the OEP mRNA is not present.

The simulations captured the expected inverse relationship between exogenous protein expression and the cell growth rate (Fig. 1A). When we made cell codon usage 60% optimal 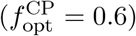, burden increased as expected when we decreased 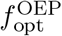 (Fig. 1A, left panel). We also ran simulations with higher levels of optimal codon usage for CP, specifically 80% and 100% optimal codon usage. Unexpectedly, we observed qualitatively different simulation behavior when we increased 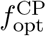 in these cases. In the most extreme case, when CP codon usage was 100% optimal (Fig. 1A, right panel), there was a complete reversal in the order of the lines, with expression of the most optimized OEP construct paradoxically creating the greatest fitness burden.

**Figure 1.**
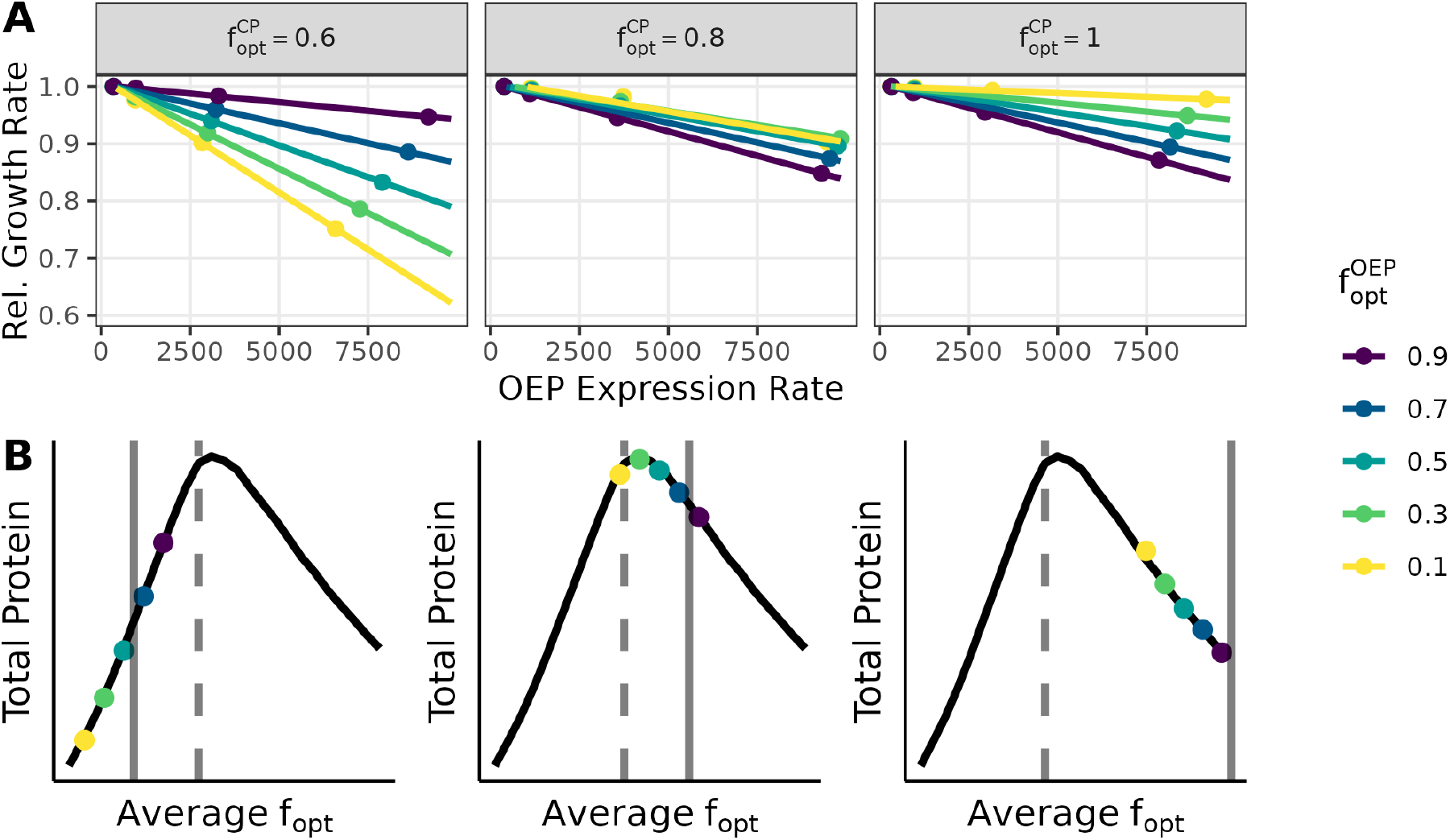
Simulation of engineered protein overexpression. (A) Simulated cell growth rate versus overexpressed protein (OEP) production rate for different fractions of optimal codon usage in the OEP 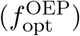 and cellular protein (CP) 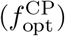. In the model, cell growth rates are the amount of CP produced over the duration of the simulation. To get the relative cell growth rate, we divide by the baseline cell growth rate when OEP is not present. OEP expression rate is the number of molecules of protein produced in a simulation divided by its duration. (B) Total protein (CP + OEP) versus average codon usage (weighted average of OEP codon usage and CP codon usage). The colored points in each plot correspond to the simulations shown in panel A, when OEP expression is maximized (highest OEP expression rate, points furthest to the right). The solid vertical line indicates 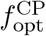, and the dashed vertical line indicates the fraction of optimal tRNA (0.7 in all cases).

This reversal can be explained by the fact that in the simulations, 100% optimal codon usage does not always maximize gene expression (Fig. 1B and Fig. 2). Specifically, when tRNA pools are partially but not completely charged, protein expression is maximized when the average *f*_opt_ of the combined set of all transcripts (for CP and OEP together) approximately matches the fraction of optimal tRNAs. (In practice, the exact location of the optimum also depends on the tRNA charging rate, see Fig. 2.) In this regime, the extent to which an overexpressed exogenous gene burdens the cell depends on the direction that expressing the OEP moves the system along the curve relative to its optimum. For example, in simulations where 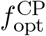 is 60% (which is slightly less than the *f*_opt_ that maximizes expression) overexpressing OEP transcripts that use fewer optimal codons than CP transcripts lowers the average *f*_opt_, moving the system away from the maximum point (Fig. 1B, left panel). As a consequence, the fraction of ribosomes sequestered on OEP transcripts increases (Fig. S1), and the cell is less able to produce its own protein. In contrast, when the codon usage of CP is 100% optimal, expressing deoptimized OEP moves the system back in the direction of the optimum (Fig. 1B, right panel). Overall, our simulations predict that burden from protein overexpression depends not only on an exogenous gene’s codon usage but also on the extent to which a cell’s codon usage matches tRNA abundances.

**Figure 2.**
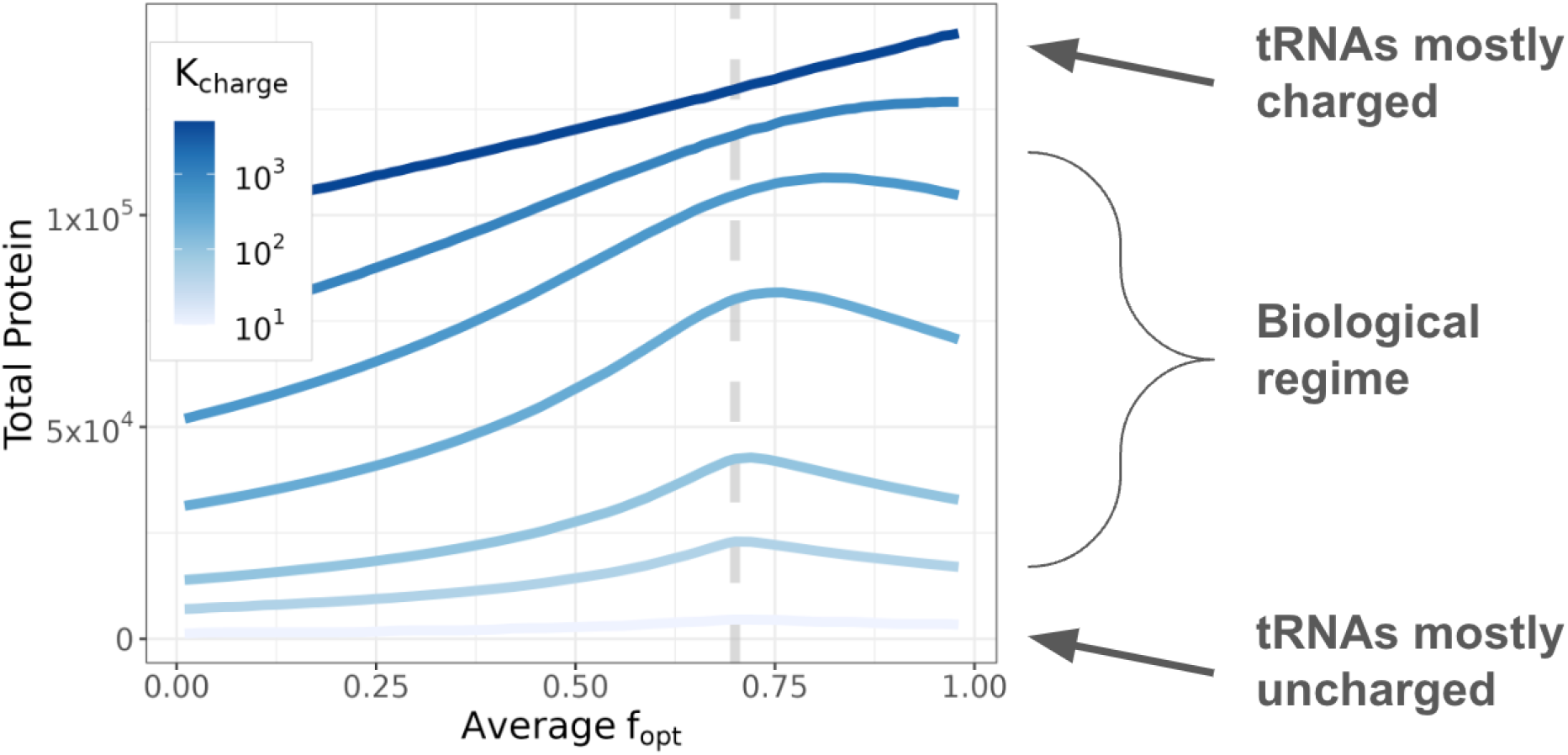
Relationship between total protein abundance and average *f*_opt_ in simulations. The dashed vertical line is the optimal tRNA fraction 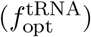, which is 0.7 in this case. When the tRNA charging rate (*K*_charge_) is moderate and tRNA pools are partially charged (biological regime) protein expression is maximized when 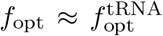. For very high tRNA charging rates, the amount of total protein produced by a cell is maximized when only optimal codons are used, but the required charging rates are much higher than is realistic for actual cells. *K*_charge_ rates are in units of s^−1^.

### 2.2 Gene expression burden depends on codon usage in *E. coli*

We tested the expectations from the model by overexpressing fluorescent proteins (FPs) modified to have different levels of codon usage bias in *E. coli*. We tested both sfGFP and mCherry2 variants. These FPs are derived from evolutionarily distant species [32] and share just 28% amino acid identity. Each FP was synthesized as a set of coding sequence (CDS) variants that used 10%, 25%, 50%, 75%, or 90% optimal codons as defined by Zhou et al. [33]. Highly expressed proteins in *E. coli* have 64% optimal codons by this metric [33, 34]. Each FP variant was cloned into a plasmid where its expression was controlled by an inducible T7 RNA polymerase promoter and one of five ribosome binding site (RBS) sequences predicted to have a range of translation initiation rates (Fig. 3A). In practice, differences in protein expression that spontaneously arose, likely due to mutations affecting T7 RNA polymerase activity, led to even more variability than the planned RBS differences between biological replicates. We used these constructs to assess how the relationship between protein overexpression and *E. coli* growth rate depended on codon usage, taking changes in fluorescence after induction during exponential growth as a measure of OEP production.

**Figure 3.**
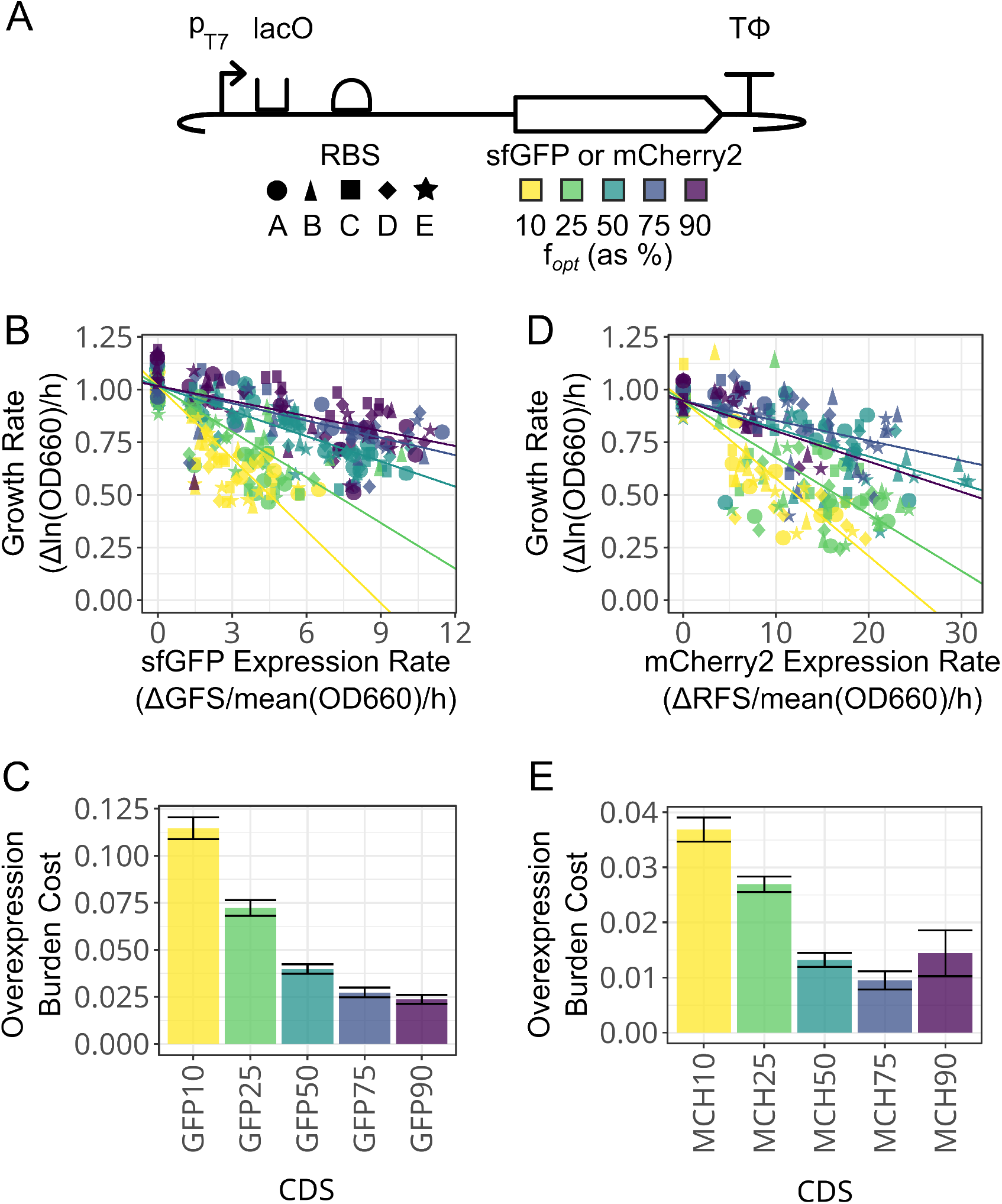
Effects of changes in codon usage on the relationship between fluorescent protein overexpression and *E. coli* growth rate. (A) Fluorescent protein constructs tested. Five different ribosome binding site (RBS) sequences were combined with five sfGFP and five mCherry2 coding sequence (CDS) variants with different percent codon optimization. The RBS and CDS shape and color legends are used in the other panels. (B) sfGFP construct rates of protein expression versus *E. coli* growth rates. Each line represents a linear regression for the respective CDS variant fit to a common *y*-intercept for no sfGFP expression. Abbreviations: GFS, green fluorescence signal; OD660, optical density at 660 nm. OD and fluorescence readings for each assay are shown in Fig. S2 and Fig. S3. (C) Overexpression burden cost for sfGFP constructs (decrease in growth rate per unit increase in the rate of protein expression) determined from the slopes of the linear regressions. Error bars are the standard error of the fits. (D) mCherry2 construct rates of protein expression versus *E. coli* growth rates. Details are as in panel B. Abbreviation: RFS, red fluorescence signal. OD and fluorescence readings for each assay are shown in Fig. S4 and Fig. S5. (E) Overexpression burden cost for mCherry2 constructs. Details are as in panel C.

**Figure 4.**
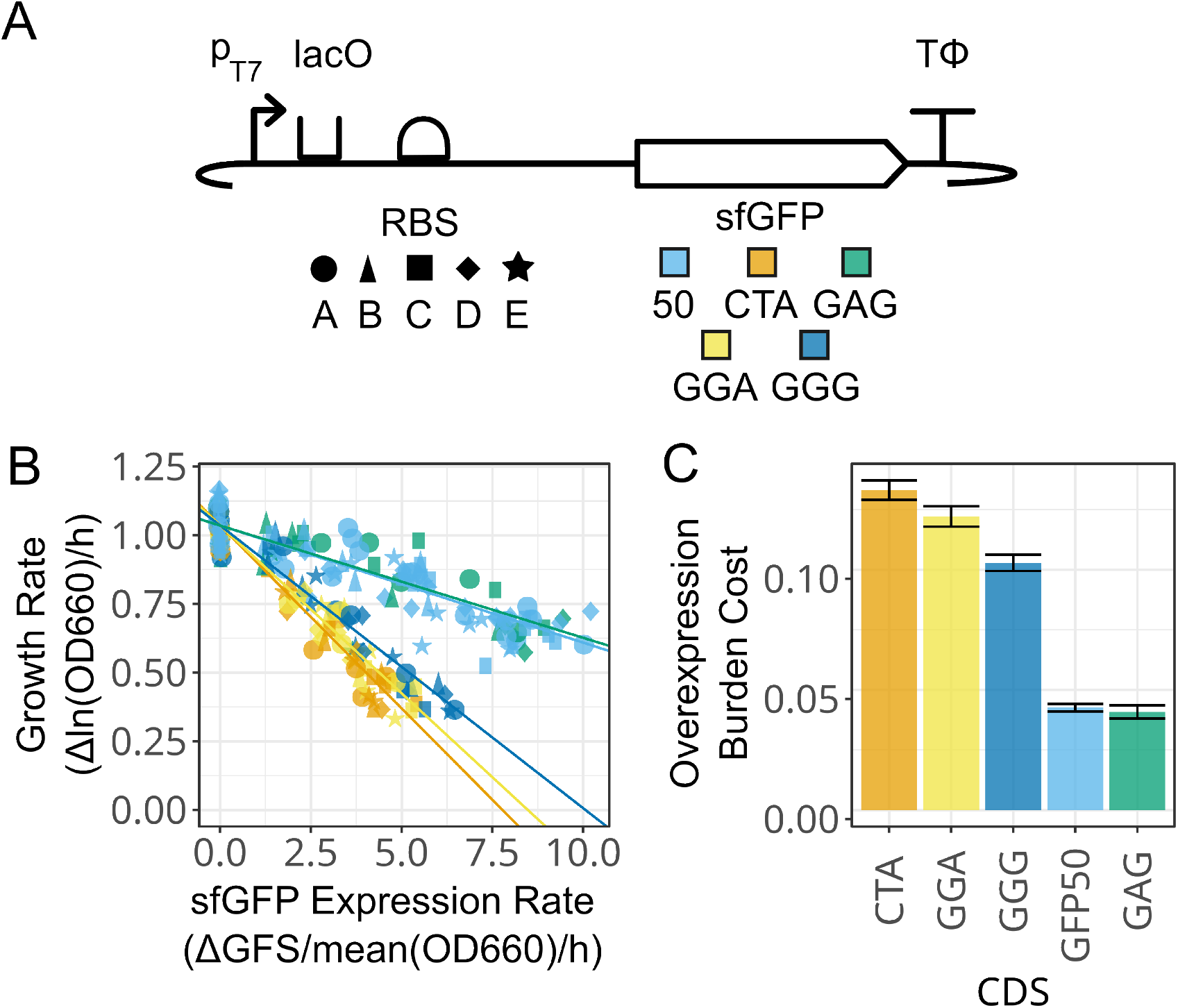
Effects of replacing codons encoding a single amino acid with a rare codon on the relationship between fluorescent protein overexpression and *E. coli* growth rate. (A) Fluorescent protein constructs tested. Five different RBS sequences were combined with the 50% optimized sfGFP CDS and four variants of this CDS shown in Table 1. The RBS and CDS shape and color legends are used in the other panels. (B) Rates of protein expression versus *E. coli* growth rates. Each line represents a linear regression for the respective CDS variant fit to a common *y*-intercept for no sfGFP expression. OD and fluorescence readings for each assay are shown in Fig. S7 and Fig. S8. (C) Overexpression burden cost for sfGFP constructs (decrease in growth rate per increase in the rate of protein expression) determined from the slopes of the linear regressions. Error bars are +/-the standard error of the fits.

Codon usage altered the relationship between how much the growth rate of *E. coli* cells slowed as they produced more sfGFP (Fig. 3B). More optimal codon usage increased the highest rate of sfGFP production achieved by a CDS variant. The 10%, 25%, 50%, and 75% optimal sfGFP constructs had maximum expression rates of 0.53, 0.73, 0.89, and 1.04 relative to the maximum expression observed for the 90% optimized variants. We also observed more burden for sfGFP variants with fewer optimal codons even though they did not achieve as high expression. CDS variants engineered with 90%, 75%, or 50% optimal codons had the smallest reductions in growth rate as the rate of protein production increased (Table S1). They reduced the growth rate by 24%, 29%, and 36%, respectively, at the highest levels of expression. Deoptimizing sfGFP to 25% or 10% optimal codons resulted in a marked increase in burden per protein produced, reducing population growth rate by 54% or 61% at the highest levels of expression. These trends can be summarized across all constructs in terms of the negative slope for the linear relationship between the two rates, which we refer to as the overexpression burden cost because it represents how much the host’s growth rate decreases for a given increase in the amount of the exogenous protein that it expresses. The mCherry2 CDSs showed a more complex trend in how overexpression burden cost varied with respect to codon usage (Fig. 3D, E). Constructs engineered with 75% and 50% optimal codons performed similarly. mCherry2 with 75% optimal codons reduced the growth rate by 22% at maximum expression, while mCherry2 at 50% optimal codons reduced the growth rate by 30% (Table S1). We observed 28% higher maximum expression with 50% optimized mCherry2 compared to 75% optimized mCherry2. mCherry2 with 25% and 10% optimal codons had more extreme impacts on *E. coli* growth rates as expression level increased. These two constructs reduced growth rates approximately 64% and 71% at maximum expression. We also found mCherry 10% to be unstable, with 29% of its replicates appearing to mutate to lose expression before they could be analyzed, which was a significantly higher fraction than that observed for the mCherry2 constructs overall (*p* = 0.0000222, binomial regression Wald test) (Fig. S6). Interestingly, the 90% optimized mCherry2 CDS was even more unstable, with 67% of constructs appearing to lose expression (*p* = 0.00000019, Wald test), and it also had the lowest maximum expression of all mCherry2 constructs. The overexpression burden cost for the 90% optimized CDS most resembled that of the 50% optimized CDS, although the difference between the overexpression burden costs of the 75% optimized and 90% optimized CDSs were not statistically significant (*p* = 0.16, likelihood ratio test). Still, these observations could indicate that the 90% optimized mCherry2 CDS is in the overoptimized regime that is predicted to exist in the simulations.

We next tested whether deoptimizing a single codon would lead to similar trends. We constructed plasmids with variants of the 50% optimized sfGFP CDS that had all of their codons for one amino acid replaced with a single rare codon for that amino acid (Table 1). We performed this targeted deoptimization for four rare codons corresponding to three amino acids (Glu, Gly, and Leu) that appear many times in the sequence of GFP. The Glu codon GAG, which has a 32% genomic frequency and is the only alternative codon to GAA for Glu, had just a 35% reduction in growth rate at maximum protein expression. This is comparable to the 37% reduction observed for the 50% optimized GFP CDS used as a baseline. Deoptimizations targeting the rarer Gly GGG, Gly GGA, and Leu CTA codons all resulted in substantial burden. They reduced growth rates at maximum expression by 63%, 61%, and 63% respectively. These results show that creating a CDS that depletes a single rare tRNA for one amino acid can also lead to high overexpression burden.

**Table 1:** Variants of the 50% optimized sfGFP CDS with all codons for a single amino acid replaced by one rare target codon.

| Amino Acid | Rare Target Codon | Genomic Frequency | Optimal to Target Replacements | Rare to Target Replacements | CDS $f_{\text{opt}}$ |
| --- | --- | --- | --- | --- | --- |
| Glu | GAG | 32% | 12 | 0 | 47.2% |
| Gly | GGA | 13% | 18 | 0 | 44.8% |
| Gly | GGG | 15% | 18 | 2 | 44.8% |
| Leu | CTA | 3% | 16 | 2 | 45.8% |

## 3 Discussion

Bacteria such as *E. coli* are commonly engineered to overexpress recombinant proteins for purification and characterization, enzymes to assimilate feedstocks and synthesize bioproducts, and biosensors and regulatory proteins to control their behavior. Here, we used stochastic simulations of gene expression to examine how codon usage influences the relationship between the rate at which a cell expresses a protein of interest and its growth rate. As expected, our simulations predict that—in many cases—replacing rare codons with optimal codons will increase protein expression and reduce the burden on a cell from producing a given amount of protein. Specifically, this is expected to be the case when a sequence begins with codon usage that is poorly matched to the cell’s preferred codon usage. However, our simulations also predict that a regime exists in which overusing optimal codons to express the protein of interest can make them become limiting and exacerbate the amount of burden from a given amount of protein production.

To test these predictions, we examined fluorescent protein genes with a range of codon usage biases expressed at different levels in *E. coli*. We found that increasing codon optimization generally led to more fluorescent protein production at less cost to the cell. We also observed elevated genetic instability at both codon usage extremes for mCherry2, including for an mCherry2 gene that seemed to exhibit signs of overoptimization. Differences in the sensitivity of the system to changes in codon usage bias for mCherry2 versus sfGFP are likely due to differences in the amino acid composition of these proteins and therefore which specific tRNA species they deplete. We further showed that deoptimizing a coding sequence by including many copies of a single rare codon for one amino acid could have as severe an effect on gene expression and burden as broadly deoptimizing its sequence to include rare codons for many amino acids.

Changing codon usage can alter other aspects of gene expression and burden that are not accounted for in our simulations. We did not alter the first 15 codons of our fluorescent protein genes to minimize well-known effects of RNA structure around the ribosome binding site and start codon on translation initiation rates [35]. RNA structures formed later in their reading frames may be highly variable between our constructs that have different codon usage. These structures could slow translation, leading to less expression and more burden [36–38]. Either of these effects of mRNA structures could explain some of the variation in expression between the constructs we tested.

It is possible that some of the burden we observe for certain constructs is due to toxic interactions between RNA and cellular components, which has been observed in similar experiments that altered synonymous codon usage in the past [39]. It is also possible that growth rates or fluorescent signals are reduced due to the formation of inclusion bodies and cell stresses related to protein misfolding, particularly for constructs that produce the most protein. Although sfGFP was evolved for improved folding and mCherry2 was evolved for lower cytotoxicity [40, 41], misfolding and toxic aggregates could arise and their levels could vary due to how codon usage and mRNA structures alter the speed of translation and affect co-translational folding [20, 42, 43]. These phenomena might introduce non-linear responses in our measurements, such that burden for overexpression increases more than expected above some threshold.

Evolutionary instability complicated our measurements [44–46]. Mutants with reduced fluorescent protein expression can arise and spread within cell populations during colony and culture growth before and after cryopreservation [5, 7]. These mutants reached a high enough frequency that they affected our experiments, even though we used a regulated promoter system to only induce fluorescent protein expression shortly before making measurements. This instability might arise if uninduced protein expression occurs and is relatively burdensome, if components of the expression system outside of the engineered plasmid (e.g., T7 RNA polymerase) are unstable and/or toxic, if induced gene expression is so burdensome that it essentially stops growth, or some combination of these and other effects. We observed heterogeneity in the fluorescent of colonies that we used to initiate different replicate measurements and a wide range of expression levels for the same RBS+CDS construct, indicating that mutants were present at appreciable frequencies in our cell stocks. We omitting replicates that exhibited very little to no fluorescence relative to other measurements of the same construct from our analyses. This type of complete evolutionary failure was especially common for the two mCherry constructs with the most extreme codon usage biases (10% and 90%),

Several studies have explored relationships between burden, gene expression, and codon usage in *E. coli*. One examined the effects of deoptimizing codons at the 3’ end of the *vioB* gene [2]. They predicted that adding rare codons would make gene expression burden more severe because slowing translation at this location would sequester more ribosomes on this mRNA, and they found this effect. Another study that compared several optimization schemes based on the codon adaptation index (CAI) also had difficulty expressing maximally codon optimized constructs [26]. A codon health index (CHI) was developed as an alternative to CAI by measuring how much use of different codons competed with native cellular proteins, as a way of coding for high expression while minimizing burden on the host cell [22]. *E. coli* cell-free transcription-translation (TX-TL) systems have been used to examine extreme codon usage while avoiding complications from evolutionary instability [22, 47].

We used Pinetree [31] to simulate dynamic charging and depletion of tRNA pools while ribosomes bind to and translate mRNAs with single-codon resolution. Pinetree has been used to simulate other bacterial gene expression processes, including transcription of mRNAs from a genome, co-transcriptional translation of mRNAs, and mRNA degradation [48, 49]. Either Pinetree or a variety of similar models that others have used to examine codon usage [2, 22, 47, 50, 51] could be extended to improve our understanding of how different bioengineering design choices affect burden. Ultimately, these simulations could refine the predictions made by design tools, such as the Operon Calculator [52] and CryptKeeper [53], so researchers can maximize protein overexpression while minimizing burden on engineered cells and reducing the incidence of evolutionary failure.

## 4 Methods

### 4.1 Gene Expression Simulation Framework

For the computational modeling, we used the Pinetree [31] gene expression simulation framework, which implements a version of the Gillespie Stochastic Simulation algorithm [54] for modeling transcription and translation dynamics at the nucleotide or codon level. In the previously published version of Pinetree, tRNAs were not explicitly simulated, and per-codon translation rates could not change dynamically during a simulation. We modified Pinetree to explicitly account for charged tRNA availability and tRNA re-charging. The updated version of Pinetree is also different from the prior version in that the per-codon translation rates depend on the availability of charged tRNA pools, which can change dynamically depending on codon usage. In the new model, pools of charged tRNAs can go to zero, if codon usage is very biased and/or the tRNA charging rate is very low. Ribosomes occupying codons with depleted tRNA pools will stall until newly charged tRNAs become available.

### 4.2 Optimal Codon Usage Model

To simulate optimal versus non-optimal codon usage in Pinetree, we used a simplified model where the total tRNA population consists of just two species—an optimal tRNA that comprises the the majority of the tRNA pool (greater than 50%), and a non-optimal tRNA that makes up the remainder of the tRNA pool. In this model, an optimal codon is a codon that corresponds to an optimal tRNA, and all transcripts are comprised of some combination of optimal and non-optimal codons. The codon-specific translation propensity (similar to the rate in an ODE model) for optimal codon translation is:

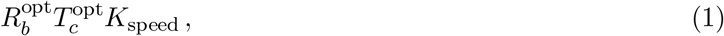

where 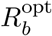 is the number of ribosomes bound to an optimal codon, 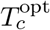 is the number of charged optimal tRNAs, and *K*_speed_ is the baseline speed of the ribosome in codons per second. Similarly, the expression for the non-optimal codon translation propensity is:

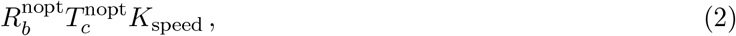

where 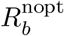 and 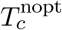 are the number of ribosomes bound to non-optimal codons and the number of non-optimal charged tRNAs, respectively. In general, 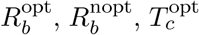, and 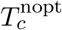 are dynamic values that can change during the simulation. In this study, these rapidly reach a steady-state within the first 10 seconds of the simulation. The simulation-specific steady-states depend on the *f*_opt_ values for OEP mRNA and CP mRNA, and the reaction rate constants for tRNA charging, ribosome binding, and ribosome elongation.

### 4.3 Simulating Exogenous Protein Overexpression

To simulate overexpression of an exogenous protein in a bacterial cell, we set up simulations with 100 identical copies of a 3000-bp mRNA transcript that represents cellular protein (CP). We then added 20 mRNA transcripts of 300-bp each that encode the overexpressed protein (OEP). We made the simplifying assumption that transcript abundances are at a steady-state and that transcription dynamics can be ignored. Each simulation begins with 500 ribosomes and 2500 total tRNAs, which are initially unbound and uncharged but quickly reach an equilibrium, where the average numbers of bound ribosomes and charged tRNAs no longer change. For the three major species in the simulation (transcripts, ribosomes, and tRNAs), we used initial total counts that are approximately 1/100th the actual abundances for each species in an *E. coli* cell during exponential growth [55, 56]. Cell growth rates and OEP production rates were calculated by taking CP and OEP abundances after 100 seconds of simulated time. Because simulations are non-deterministic, for every individual simulation we ran 3 replicates and used protein abundances averaged over these replicates. To get steady-state charged tRNA and steady-state bound ribosomes amounts, we took an average over the last 50 seconds of the simulation run, after it had reached equilibrium.

### 4.4 Calibration of Simulation Parameters

The major free parameters in these simulations are the rate constants for tRNA charging (*K*_charge_), ribosome binding to cell RBS (*K*_bind_), and ribosome elongation (*K*_speed_). To calibrate these parameters, we manually fit ribosome and tRNA steady-states in the baseline simulation (when OEP is not present) to empirical measurements. In exponentially growing *E. coli* cells, at least 80% of the total ribosome content is bound to mRNA [57], and 75% of the tRNA pool is charged with an amino acid [58]. Using these values as our target, we set *K*_bind_ to 1 *×* 10^7^ M^−1^s^−1^ (which we had used in prior simulations [48]) and then chose values for *K*_charge_ and *K*_speed_ that achieved close to the target steady-state quantities. Figure S9 shows the entire grid search over this parameter space. Finally, for 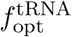, we chose a value of 0.7, which means that 70% of the total tRNA pool in the simulation is optimal tRNA and 30% is non-optimal tRNA. (Note that the fractions of *charged* tRNAs can and will differ from these values, depending on the codon usage of the CP and OEP and various simulation parameters.) A summary of all simulation parameters and their values is provide in Table 2.

**Table 2:** Parameters for overexpression simulations.

| Parameter | Value(s) | Units |
| --- | --- | --- |
| $K_{\text{speed}}$ | 2 | codons/s |
| $K_{\text{charge}}$ | 100 | $\text{s}^{-1}$ |
| $K_{\text{bind}}$ (CP RBS) | $1 \times 10^7$ | $\text{M}^{-1}\text{s}^{-1}$ |
| $K_{\text{bind}}$ (OEP RBS) | $1 \times 10^6 - 1 \times 10^8$ | $\text{M}^{-1}\text{s}^{-1}$ |
| $f_{\text{opt}}^{\text{CP}}$ | 0.6, 0.8, 1.0 | - |
| $f_{\text{opt}}^{\text{OEP}}$ | 0.1–0.9 | - |
| $f_{\text{opt}}^{\text{tRNA}}$ | 0.7 | - |

### 4.5 Plasmid Design and Assembly

DNA sequences encoding mCherry2 and sfGFP CDSs with different codon usage biases were generated using a Python script that randomly replaced codons between optimal and nonoptimal possibilities until the desired level of optimization was reached. The first 15 codons were left unchanged to reduce the chances of substantially altering translation initiation rates [35]. Additionally, we mutated mCherry2 to eliminate an alternative start codon [59]. Optimal codon classifications were from a previous analysis of highly expressed *E. coli* genes [33]. For sfGFP genes in which we deoptimized all codons for one amino acid, 2-3 codons for other amino acids had to be mutated from optimal to rare codons to make the sequence compatible with DNA synthesis. Primers and plasmids are shown in Tables S2 & S3. The full coding sequences tested are provided in Tables S4, S5, & S6.

We initially cloned these CDS constructs into plasmid pET21 and performed experiments using variable concentrations of IPTG to achieve different expression levels, but this resulted in differences in the timing of induction that complicated data analysis [60]. Therefore, we designed plasmids in which we cloned new RBS sequences predicted to have 0.25 (A), 0.5 (B), 2 (C), or 4 (D) times the strength of the pET21 vector RBS (E). Alternative ribosome binding sites were generated *in silico* by introducing random mutations and predicting translation initiation rates using OSTIR [61] until they matched the desired strength in the context of a complete sfGFP expression plasmid.

CDS variants were ordered as gBlocks (Integrated DNA Technologies). Golden Gate assembly overhangs and alternative RBS sequences were either included in the gBlocks or added in PCR primers. Plasmid pET21b(+) was domesticated to remove three BsmBI cut sites using PaqCI Golden Gate assembly (GGA). Parts were first assembled into an insulated part plasmid [62] using BsmBI GGA and chemically transformed into *E. coli* NEB 5-alpha cells (New England Biolabs). Part plasmids and the pET21b(+) backbone were then assembled into complete expression constructs using PaqCI GGA, chemically transformed into NEB 5-alpha, and subcloned into *E. coli* Tuner(DE3) cells (Novagen). Plasmids were confirmed to have the designed sequence and to be monomeric using nanopore sequencing (Plasmidsaurus).

### 4.6 Fluorescent Protein Overexpression Burden Assays

*E. coli* strains with FP overexpression plasmids were cultured in LB supplemented with 100 µg/ml carbenicillin (Crb). Strains were struck from –80°C glycerol stocks onto LB + Crb plates and allowed to grow overnight at 30 °C or 37 °C. The next day, single colonies were picked at random and allowed to grow overnight in 5 mL LB + Crb at 30 °C or 37 °C with orbital shaking at 200 RPM over a 1-inch diameter. 10 µL of overnight culture was added to 190 µL of fresh LB + Crb in a 96-well microplate and allowed to precondition in a Tecan Infinite Pro M200 microplate reader at 37 °C with orbital shaking over a 1-mm radius at 432 RPM for 11 min 40 sec per 15-min measurement cycle for a total of 2 to 2.5 hr. Then, cultures were normalized to an OD660 reading of approximately 0.2 and induced by adding 0.19 mM isopropyl *β*-D-1-thiogalactopyranoside (IPTG) inducer using an automated liquid handler or 0 mM to measure uninduced growth rate. This inducer concentration was optimized by conducting preliminary tests of the 50% optimal GFP constructs with 0.07 mM to 0.25 mM IPTG and selecting a concentration that produced a high dynamic range of growth rates. To control for position effects in the plate, cells with different constructs were randomly distributed across wells, excluding the perimeter. Cells were then induced and immediately returned to the microplate reader. OD660 and fluorescence measurements were taken every 15 minutes. sfGFP was measured at 480 nm excitation and 525 nm emission wavelengths. mCherry2 was measured at 590 nm excitation and 645 nm emission wavelengths.

Following an approach used in prior studies [2, 63], *E. coli* growth rate and FP expression rates were analyzed in 1-hour sliding windows. Growth rate was defined as 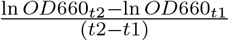. FP expression rate was defined as 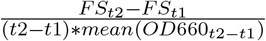, where FS is the fluorescent signal from sfGFP or mCherry2. The starting time for windows used to fit growth rates was 2 h 15 min post-induction for both GFP and mCherry2. FP expression rates were fit from time windows beginning 2 h 15 min post-induction for sfGFP variants and 3 h 15 min post-induction for mCherry2 variants. These times were selected to correspond to high growth rate variability and maximum fluorescent protein production rates. The delay in measuring mCherry2 expression reflects its slower maturation time.

For analyzing the relationship between growth and expression rates, linear regression was performed with a common *y*-intercept (zero expression) within each group of CDS variants (percent optimized sfGFP, percent optimized mCherry2, or single-codon deoptimized sfGFP). Induced constructs with less than 10% of the maximal normalized fluorescence were presumed to have completely mutated by the time of the expression measurement and were dropped from the analysis (Fig. S6). The mean of the top 10% observed expression values for each CDS variant was used for reporting growth rates at maximal expression. The final parameters fit for each construct are provided in Table S1.

## 5 Data and Code Availability

Code and source data for the simulations and experiments in this study are available in Zenodo (https://doi.org/10.5281/zenodo.14232327).

## 6 Author Contributions

Cameron T. Roots: Conceptualization; Software; Formal Analysis; Investigation; Visualization; Methodology; Writing—original draft; Writing—review and editing. Alexis M. Hill: Software; Formal Analysis; Investigation; Visualization; Methodology; Writing—original draft; Writing—review and editing. Claus O. Wilke: Conceptualization; Formal Analysis; Supervision; Methodology; Funding acquisition; Writing— review and editing. Jeffrey E. Barrick: Conceptualization; Formal Analysis; Supervision; Funding acquisition; Writing—original draft; Writing—review and editing.

## 8 Acknowledgements

The authors acknowledge the Texas Advanced Computing Center (TACC) at The University of Texas at Austin for providing high-performance computing resources. This work was supported by the National Institutes of Health (R01GM088344 to COW and JEB) and the National Science Foundation (MCB-2123996 to JEB). COW also acknowledges support from the Blumberg Centennial Professorship in Molecular Evolution.

