## Supporting Information for "Modeling and measuring how codon usage modulates the relationship between burden and yield during protein overexpression in bacteria"

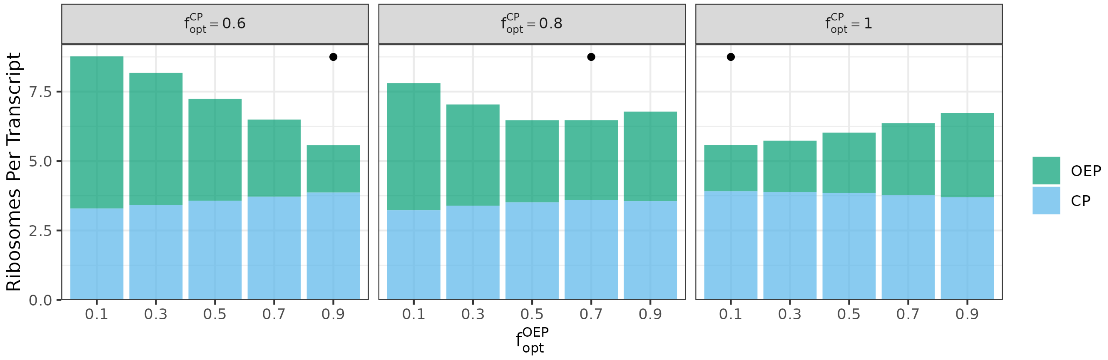

Supplementary Figure S1: **Ribosomes per transcript in overexpression simulations.** Data is for simulations in which the rate of OEP translation initiation is 3 times that of CP initiation. Points indicate the optimal codon fraction in OEP ($f_{\text{opt}}^{\text{OEP}}$) that minimizes cell burden.

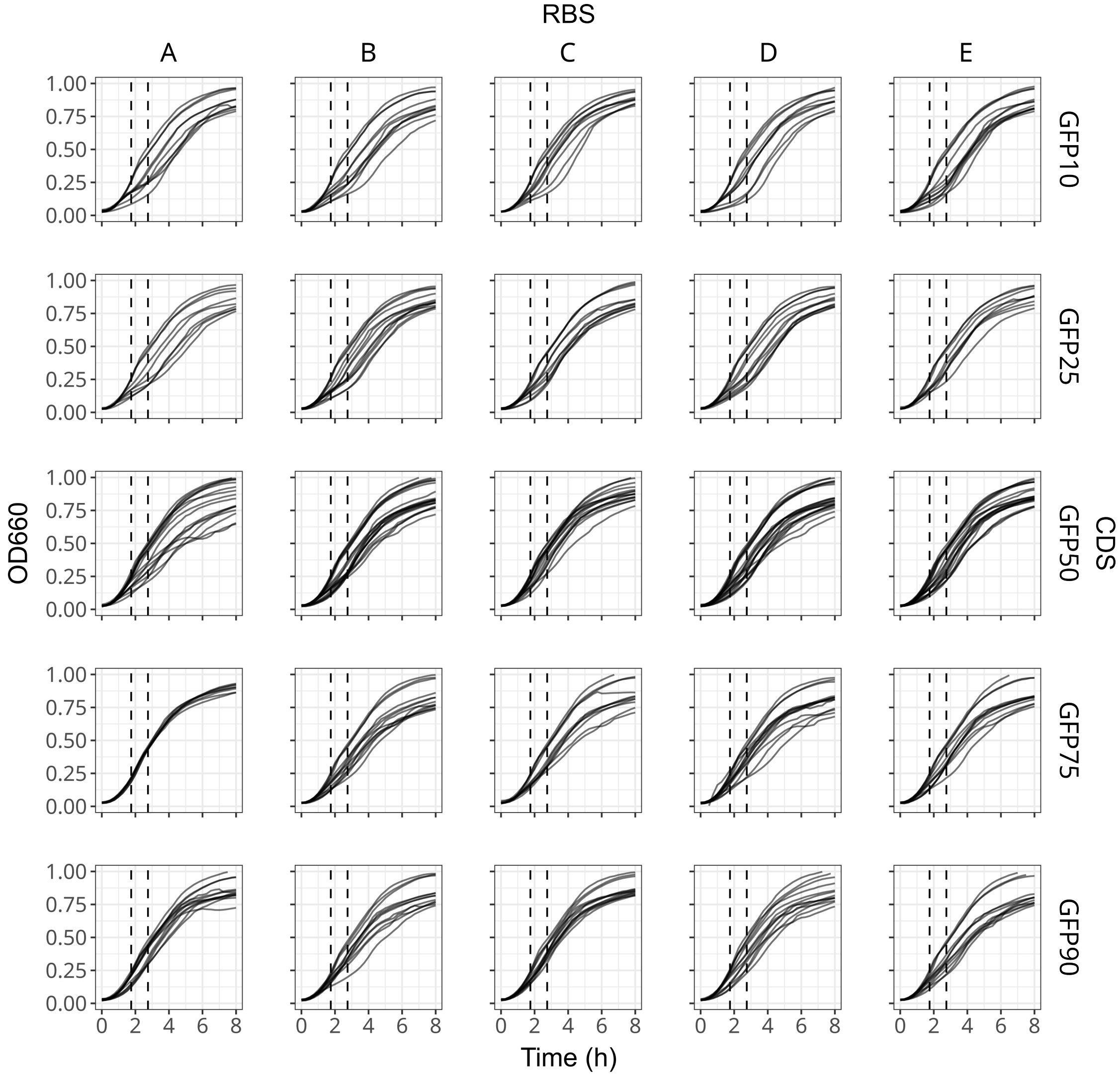

Supplementary Figure S2: **Growth curves of sfGFP constructs engineered to a certain $f_{\mathbf{opt}}$. Vertical dashed lines indicate the window used for fitting the growth rate.**

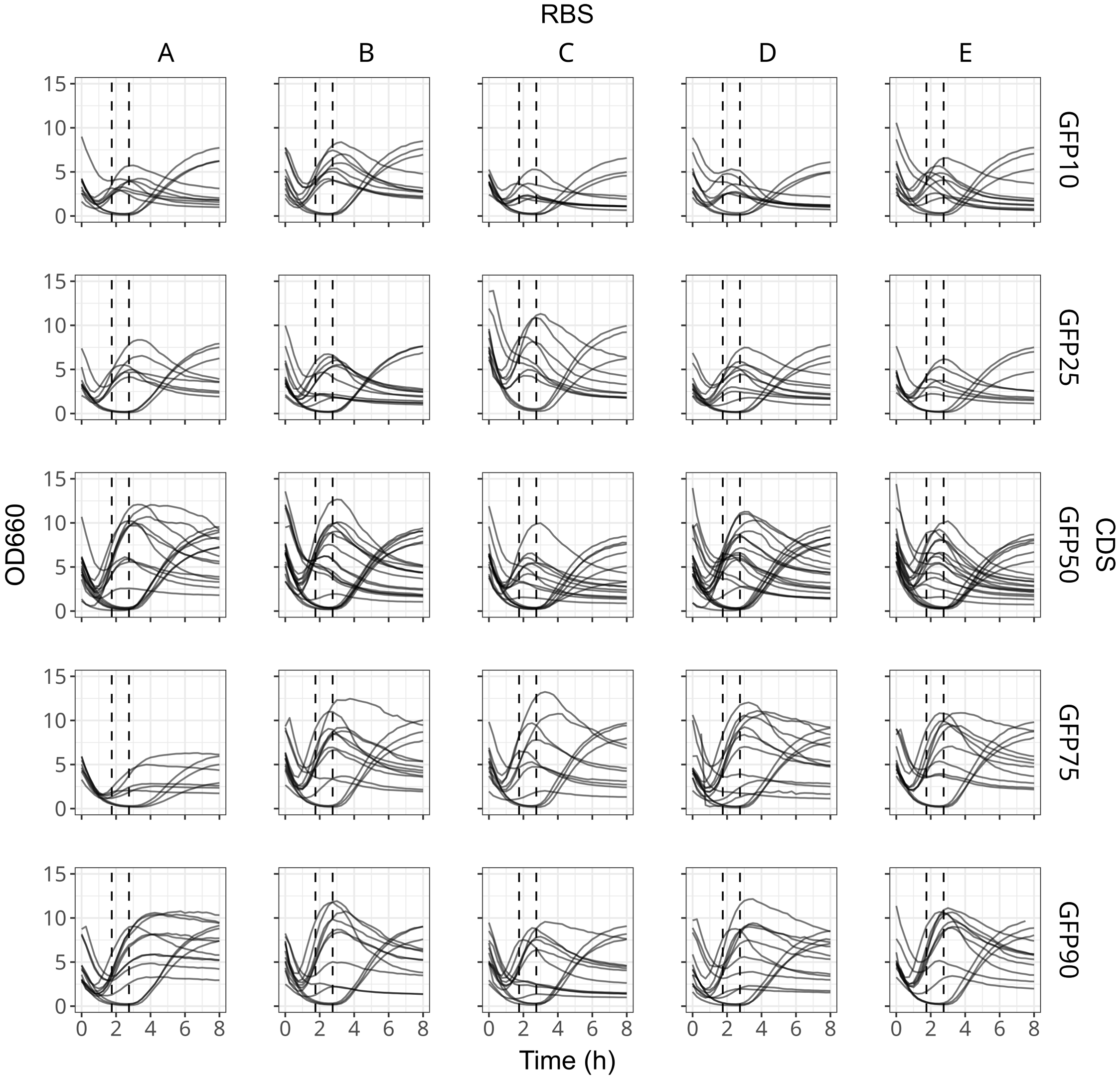

Supplementary Figure S3: **Fluorescence expression curves of sfGFP constructs engineered to a certain $f_{\mathbf{opt}}$.** Vertical dashed lines indicate the window used for fitting the fluorescent protein expression rate.

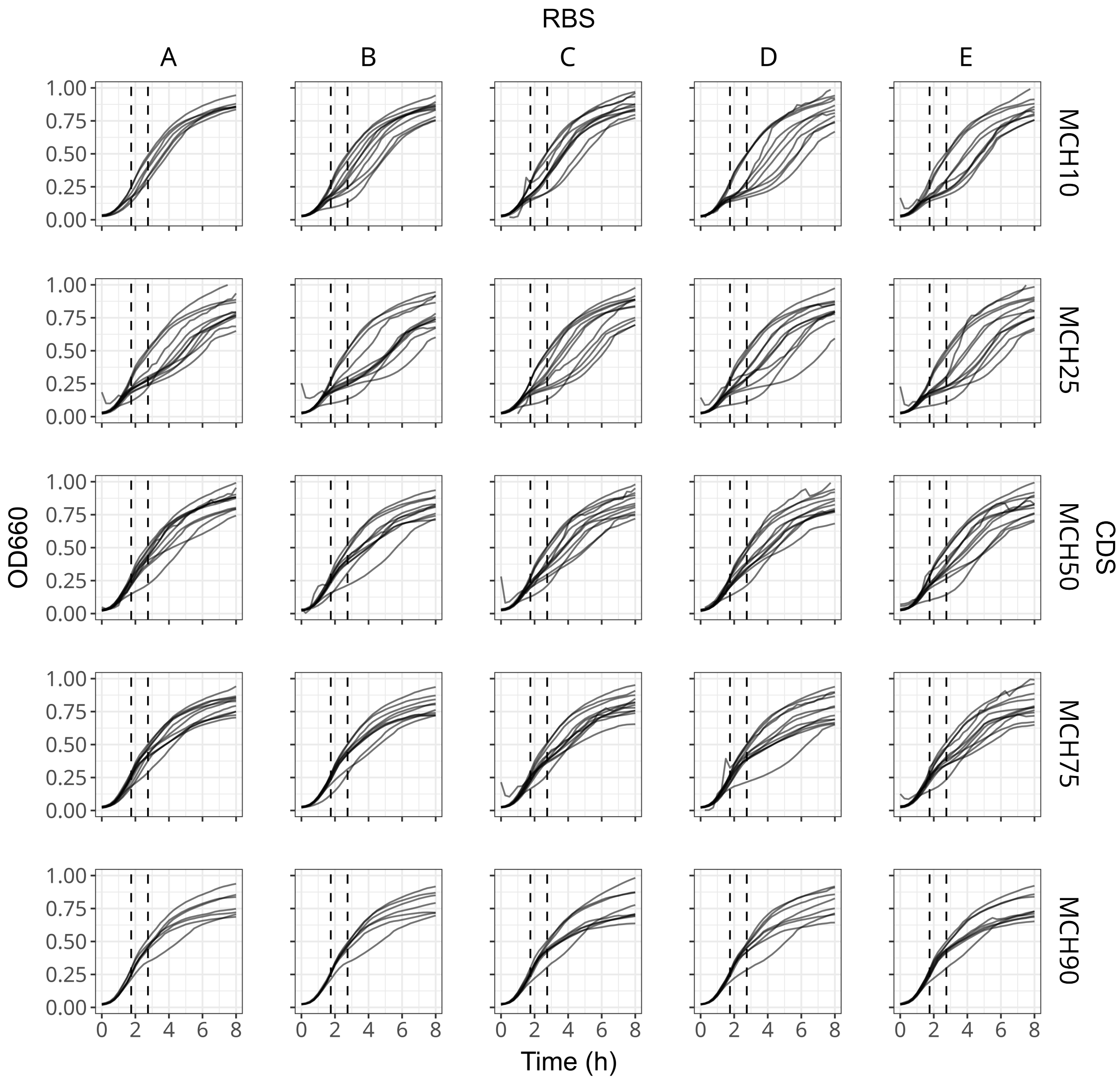

Supplementary Figure S4: **Growth curves of mCherry2 constructs engineered to a certain $f_{\mathbf{opt}}$.** Vertical dashed lines indicate the window used for fitting the growth rate.

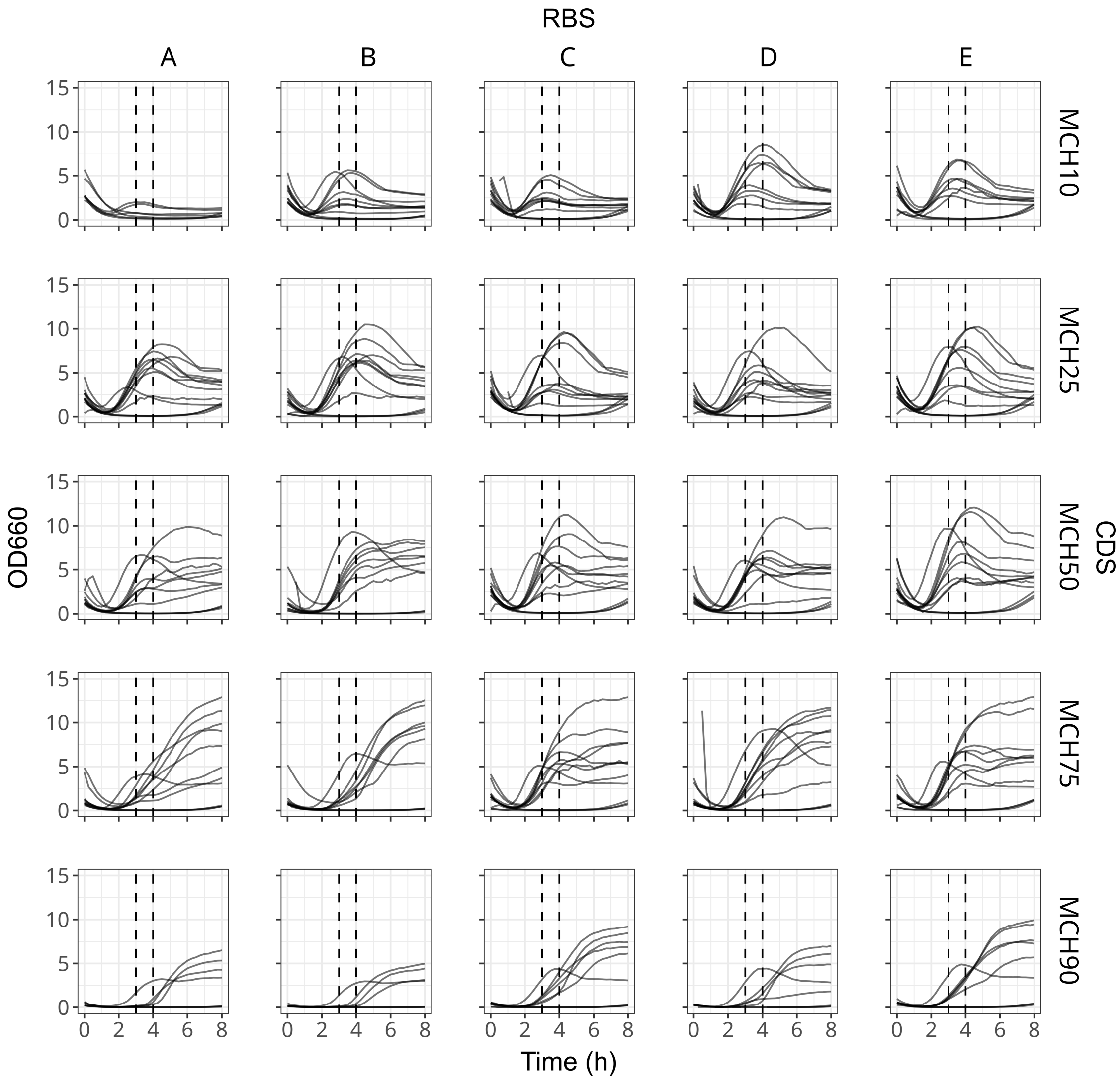

Supplementary Figure S5: **Fluorescence expression curves of mCherry2 constructs engineered to a certain $f_{\mathbf{opt}}$.** Vertical dashed lines indicate the window used for fitting the fluorescent protein expression rate.

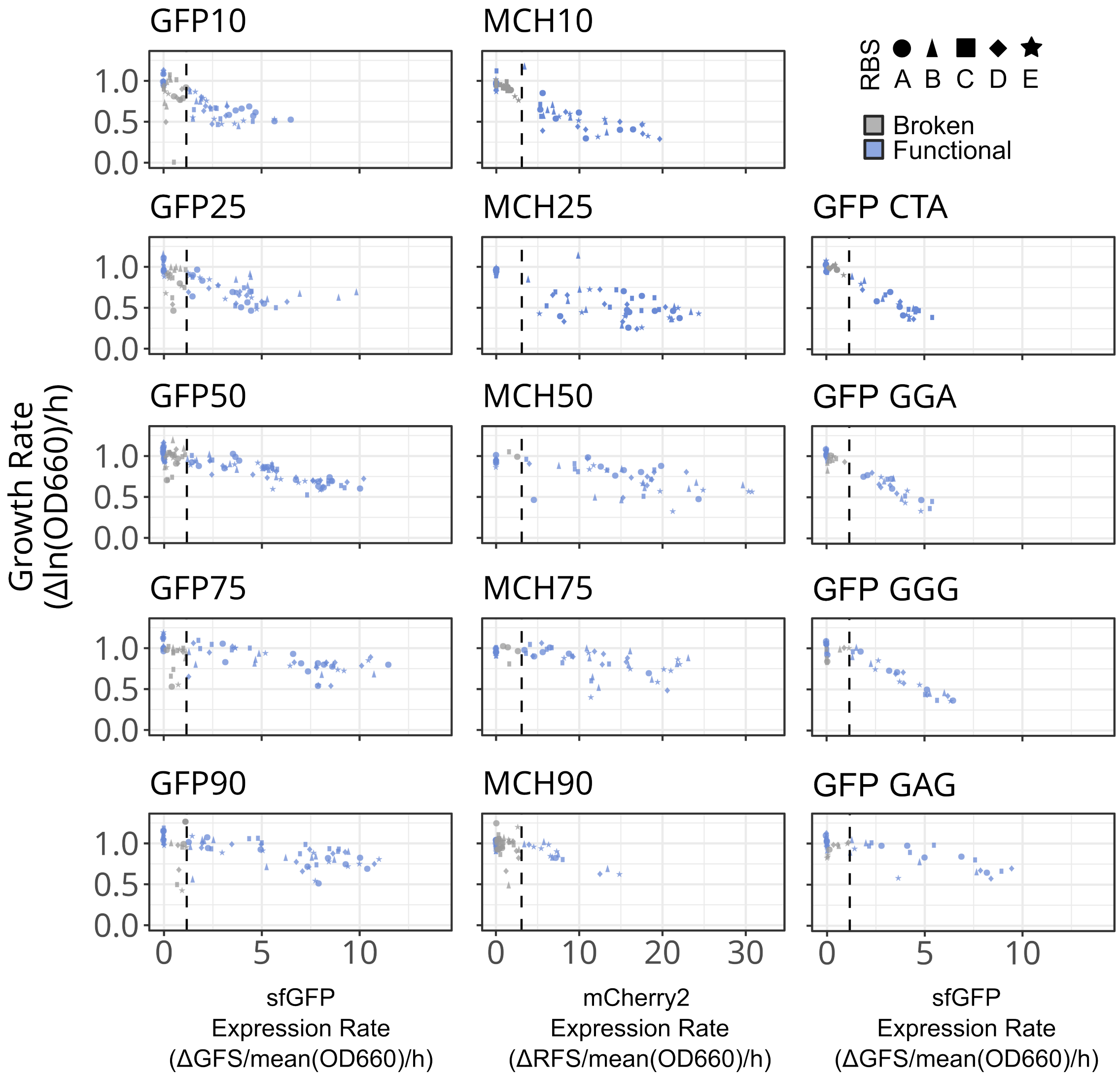

Supplementary Figure S6: **Rates of protein expression versus *E. coli* growth rates for each CDS.** Vertical line represents 10% of the observed maximum expression for each fluorescent protein. Induced constructs below this threshold are presumed to be broken by the time of measurement.

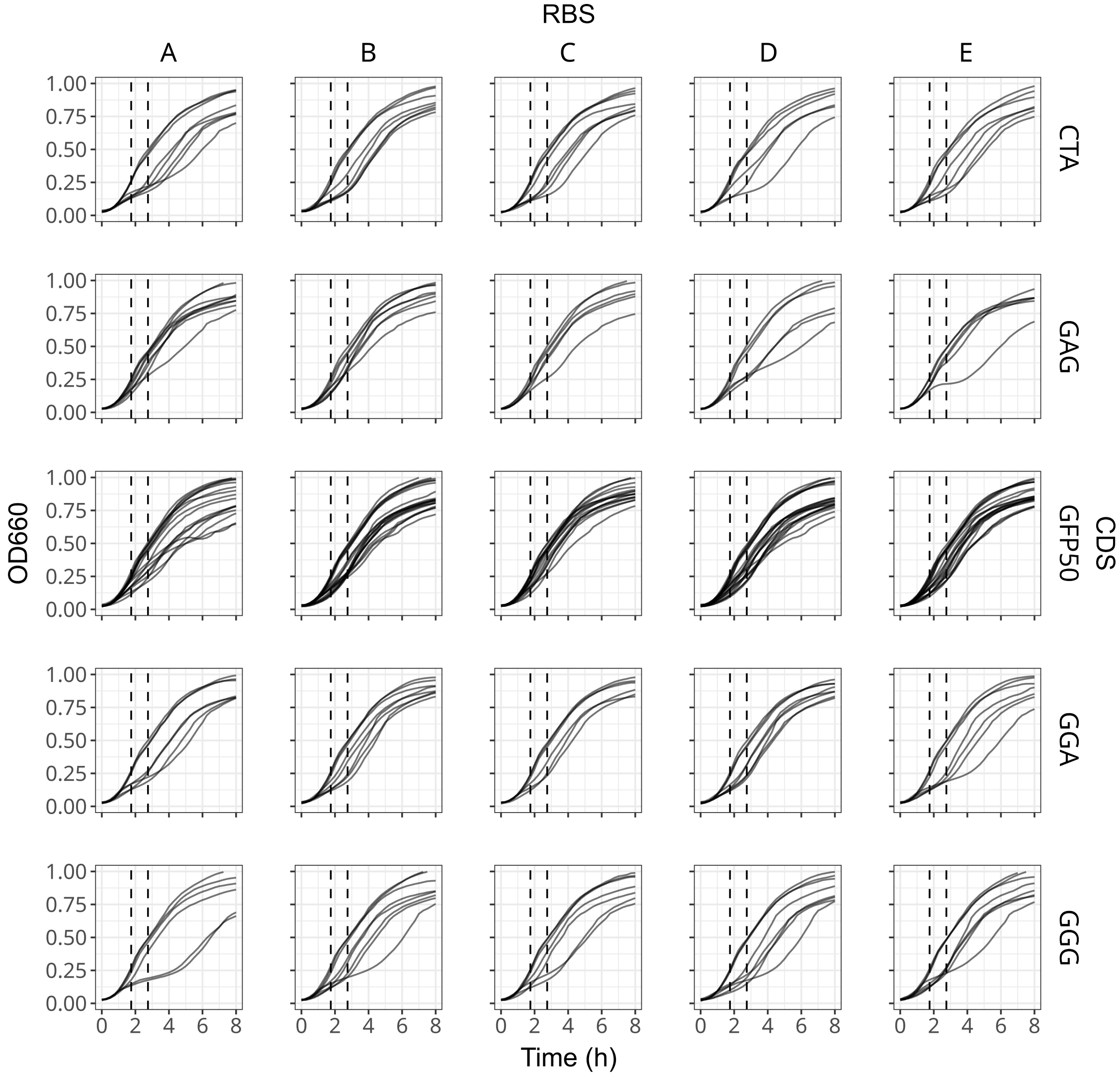

Supplementary Figure S7: **Growth curves of sfGFP constructs where one amino acid was maximally encoded by a rare codon.** Vertical dashed lines indicate the window used for fitting the growth rate.

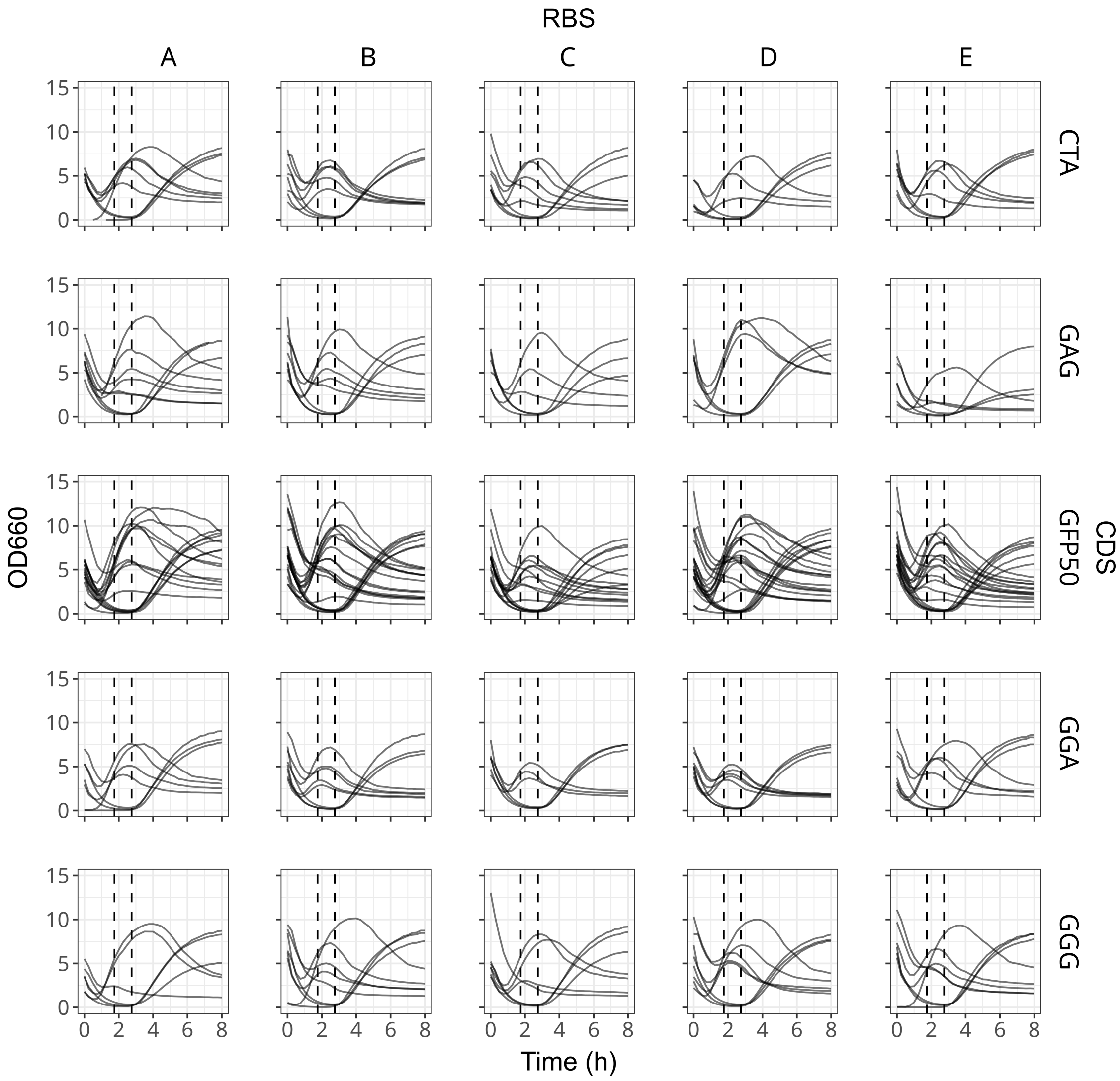

Supplementary Figure S8: **Fluorescence expression curves of codon saturated sfGFP constructs where one amino acid was maximally encoded by a rare codon.** Vertical dashed lines indicate the window used fitting the fluorescent protein expression rate.

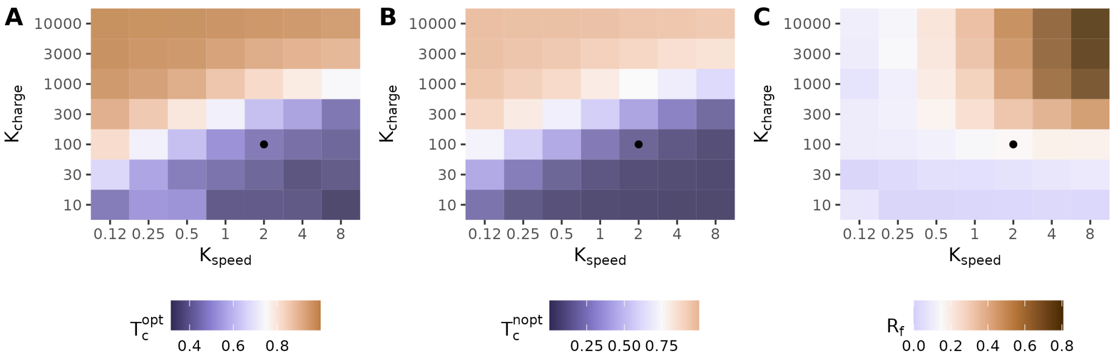

Supplementary Figure S9: **Calibration of tRNA and ribosome steady-states.** Steady-state levels of charged optimal tRNA (A), charged non-optimal tRNA (B), and free (unbound) ribosomes (C) for different charging rate ($K_{\text{charge}}$) and ribosome speed ($K_{\text{speed}}$) parameter values. Steady-states quantities are given as fractions (i.e., as the numbers of charged tRNA molecules or unbound ribosomes divided by total amounts for each species in the simulation). Color scale midpoints are the steady-states observed in real cells growing exponentially, where tRNAs are about 75% charged, and ribosomes are about 20% unbound. Overexpression simulations use a ribosome speed of 2 and a charging rate of 100; the corresponding tiles here have been annotated with black points.

Supplementary Table S1: **Growth rate and expression parameters fit for all constructs**

| CDS | Maximum Expression Rate ($\Delta$FS/mean(OD660)/h) | Growth Rate at Maximum Expression ($\Delta$ln(OD660)/h) | Fit Intercept | Fit Slope ($-$Overexpression Burden Cost) | Calculated Burden at Maximum Expression |
|---|---|---|---|---|---|
| GFP10 | 5.42 | 0.55 | 1.02 | -0.115 | 61% |
| GFP25 | 7.54 | 0.619 | 1.02 | -0.072 | 54% |
| GFP50 | 9.10 | 0.68 | 1.02 | -0.040 | 36% |
| GFP75 | 10.66 | 0.81 | 1.02 | -0.027 | 29% |
| GFP90 | 10.26 | 0.79 | 1.02 | -0.024 | 24% |
| MCH10 | 18.27 | 0.37 | 0.95 | -0.037 | 71% |
| MCH25 | 22.38 | 0.46 | 0.95 | -0.027 | 64% |
| MCH50 | 27.98 | 0.58 | 0.95 | -0.013 | 30% |
| MCH75 | 21.94 | 0.83 | 0.95 | -0.009 | 22% |
| MCH90 | 13.58 | 0.66 | 0.95 | -0.014 | 21% |
| CTA | 4.90 | 0.45 | 1.04 | -0.133 | 63% |
| GAG | 8.92 | 0.65 | 1.04 | -0.041 | 35% |
| GGA | 5.17 | 0.43 | 1.04 | -0.122 | 61% |
| GGG | 6.33 | 0.39 | 1.04 | -0.102 | 63% |

### Supplementary Table S2: **Primers used in this study**

| Name | Purpose | Sequence |
|---|---|---|
| T7tag_F | Remove T7 tag from pET21b(+) | CATATGTATATCTCCTTCTTAAAGTTAAACAAAATTATTTCTAGAGGGGAATTGTTATCC |
| T7tag_R | Remove T7 tag from pET21b(+) | GATCCGAATTCGAGCTCCGT |
| Histag_F | Remove His tag from pET21b(+) | TTATTATTACTCGAGTGCGGCCGCAA |
| Histag_R | Remove His tag from pET21b(+) | TGAGATCCGGCTGCTAACAAAGCC |
| pET21_F1 | Make PaqCI domesticated pET21 Dropout Vector | TTACGCGGTCTCGAGCCACGCTCACCGGCTCCAG |
| pET21_R1 | Make PaqCI domesticated pET21 Dropout Vector | TTACGCGGTCTCGACAAAGCCCGAAAGGAAGCTGAGTTGG |
| pET21_F2 | Make PaqCI domesticated pET21 Dropout Vector | TTACGCGGTCTCGACGCCGGACGCATCGTGGCCG |
| pET21_R2 | Make PaqCI domesticated pET21 Dropout Vector | TTACGCGGTCTCGGGCTCTCGCGGAATCATTGCAGCACTG |
| pBTK1001_F | Make PaqCI domesticated pET21 Dropout Vector | TTACGCGGTCTCGTTGTTATTACGCAGGTGGAAAGTGAAACGTG |
| pBTK1001_R | Make PaqCI domesticated pET21 Dropout Vector | TTACGCGGTCTCGGCGTATCCAGCAGGTGTATAAACGCAG |
| GFP_F | Assembling GFP Parts | TTACGCGGTCTCGTTGTTATTACGCAGGTGGAAAGTGAAACGTGATTTCATG |
| GFP_R | Assembling GFP Parts | TTACGCGGTCTCGGCGTATCCAGCAGGTGTATAAACGCAGAAAGGCC |
| MCH_F | Assemblying mCherry Parts | ATGCCGTCTCATCGGCACCTGCTATACATATGGTGAGCAAGGGCGAG |
| MCH_R | Assembling mCherry Parts | GCATCGTCTCAGGTCACCTGCCTTTGTTAGCAGCCGGATCTCATTATTA |
| RBS.A_F | Assembling RBS.A Part | GCATCGTCTCAGGTCACCTGCACCATATGTATAACTCCTTCTTAAAGTTAAACAAAAT |
| RBS.B_F | Assembling RBS.B Part | GCATCGTCTCAGGTCACCTGCACCATATGTATATATCCTTCTTAAAGTTAAACAAAAT |
| RBS.C_F | Assembling RBS.C Part | GCATCGTCTCAGGTCACCTGCACCATATGTATATCTCCTTCTTAAAGTTAAATAAAAT |
| RBS.D_F | Assembling RBS.D Part | GCATCGTCTCAGGTCACCTGCACCATATGTATATCTCCTCCTTAAAGTTAAACAAAAT |
| RBS.E_F | Assembling RBS.E Part | GCATCGTCTCAGGTCACCTGCACCATATGTATATCTCCTTCTTAAAGTTAAACAAAAT |
| RBS_R | Assembling RBS Parts | ATGCCGTCTCATCGGCACCTGCCCGGCGTAGAGGATCGAGATC |

Supplementary Table S3: **Plasmids Used in this study**

| Plasmid | Source | Plasmid | Source |
|---|---|---|---|
| pET21b(+) | Novagen | pET21-Domesticated | This study |
| pBTK1001 | Lariviere 2024 | | |
| pBTK1001-sfGFP.10 | This study | pET21-RBS.A-sfGFP.GGG | This study |
| pBTK1001-sfGFP.25 | This study | pET21-RBS.B-sfGFP.GGG | This study |
| pBTK1001-sfGFP.50 | This study | pET21-RBS.C-sfGFP.GGG | This study |
| pBTK1001-sfGFP.75 | This study | pET21-RBS.D-sfGFP.GGG | This study |
| pBTK1001-sfGFP.90 | This study | pET21-RBS.E-sfGFP.GGG | This study |
| pBTK1001-sfGFP.GGG | This study | pET21-RBS.A-sfGFP.GGA | This study |
| pBTK1001-sfGFP.GGA | This study | pET21-RBS.B-sfGFP.GGA | This study |
| pBTK1001-sfGFP.GAG | This study | pET21-RBS.C-sfGFP.GGA | This study |
| pBTK1001-sfGFP.CTA | This study | pET21-RBS.D-sfGFP.GGA | This study |
| | | pET21-RBS.E-sfGFP.GGA | This study |
| pBTK1001-mCherry2.10 | This study | pET21-RBS.A-sfGFP.GAG | This study |
| pBTK1001-mCherry2.25 | This study | pET21-RBS.B-sfGFP.GAG | This study |
| pBTK1001-mCherry2.50 | This study | pET21-RBS.C-sfGFP.GAG | This study |
| pBTK1001-mCherry2.75 | This study | pET21-RBS.D-sfGFP.GAG | This study |
| pBTK1001-mCherry2.90 | This study | pET21-RBS.E-sfGFP.GAG | This study |
| pBTK1001-RBS.A | This study | pET21-RBS.A-sfGFP.CTA | This study |
| pBTK1001-RBS.B | This study | pET21-RBS.B-sfGFP.CTA | This study |
| pBTK1001-RBS.C | This study | pET21-RBS.C-sfGFP.CTA | This study |
| pBTK1001-RBS.D | This study | pET21-RBS.D-sfGFP.CTA | This study |
| pBTK1001-RBS.E | This study | pET21-RBS.E-sfGFP.CTA | This study |
| pET21-RBS.A-sfGFP.10 | This study | pET21-RBS.A-mCherry2.10 | This study |
| pET21-RBS.B-sfGFP.10 | This study | pET21-RBS.B-mCherry2.10 | This study |
| pET21-RBS.C-sfGFP.10 | This study | pET21-RBS.C-mCherry2.10 | This study |
| pET21-RBS.D-sfGFP.10 | This study | pET21-RBS.D-mCherry2.10 | This study |
| pET21-RBS.E-sfGFP.10 | This study | pET21-RBS.E-mCherry2.10 | This study |
| pET21-RBS.A-sfGFP.25 | This study | pET21-RBS.A-mCherry2.25 | This study |
| pET21-RBS.B-sfGFP.25 | This study | pET21-RBS.B-mCherry2.25 | This study |
| pET21-RBS.C-sfGFP.25 | This study | pET21-RBS.C-mCherry2.25 | This study |
| pET21-RBS.D-sfGFP.25 | This study | pET21-RBS.D-mCherry2.25 | This study |
| pET21-RBS.E-sfGFP.25 | This study | pET21-RBS.E-mCherry2.25 | This study |
| pET21-RBS.A-sfGFP.50 | This study | pET21-RBS.A-mCherry2.50 | This study |
| pET21-RBS.B-sfGFP.50 | This study | pET21-RBS.B-mCherry2.50 | This study |
| pET21-RBS.C-sfGFP.50 | This study | pET21-RBS.C-mCherry2.50 | This study |
| pET21-RBS.D-sfGFP.50 | This study | pET21-RBS.D-mCherry2.50 | This study |
| pET21-RBS.E-sfGFP.50 | This study | pET21-RBS.E-mCherry2.50 | This study |
| pET21-RBS.A-sfGFP.75 | This study | pET21-RBS.A-mCherry2.75 | This study |
| pET21-RBS.B-sfGFP.75 | This study | pET21-RBS.B-mCherry2.75 | This study |
| pET21-RBS.C-sfGFP.75 | This study | pET21-RBS.C-mCherry2.75 | This study |
| pET21-RBS.D-sfGFP.75 | This study | pET21-RBS.D-mCherry2.75 | This study |
| pET21-RBS.E-sfGFP.75 | This study | pET21-RBS.E-mCherry2.75 | This study |
| pET21-RBS.A-sfGFP.90 | This study | pET21-RBS.A-mCherry2.90 | This study |
| pET21-RBS.B-sfGFP.90 | This study | pET21-RBS.B-mCherry2.90 | This study |
| pET21-RBS.C-sfGFP.90 | This study | pET21-RBS.C-mCherry2.90 | This study |
| pET21-RBS.D-sfGFP.90 | This study | pET21-RBS.D-mCherry2.90 | This study |
| pET21-RBS.E-sfGFP.90 | This study | pET21-RBS.E-mCherry2.90 | This study |

Supplementary Table S4: **Sequences of sfGFP genes with different codon usage**

| CDS Name | Nucleotide Sequence |
| --- | --- |
| GFP10 | ATGAGCAAAGGTGAAGAACTGTTTACCGGCGTTGTGCCGATTCTGGTGGAGCTAGATGGGGATGTGAAT |
| | GGACATAAATTTAGCGTGCGGGGGGAGGGAGAAGGGGATGCGACGAATGGGAAACTTACATTAAAATT |
| | TATTTGTACAACGGGAAAACTCCCTGTCCCATGGCCTACATTGGTGACGACGCTAACATATGGAGTCCA |
| | ATGTTTTAGCCGGTATCCCGATCATATGAAACGACATGATTTCTTTAAAAGCGCGATGCCCGAGGGATA |
| | TGTGCAAGAGAGGACCATTAGCTTTAAAGATGATGGAACATATAAAACACGGGCGGAAGTTAAATTTGA |
| | GGGGGATACGCTAGTGAATAGAATTGAGCTAAAAGGTATTGATTTTAAAGAGGATGGGAATATTCTTGG |
| | ACATAAACTAGAGTATAATTTCAACAGCCATAATGTGTATATTACAGCCGATAAACAAAAAAATGGGAT |
| | AAAAGCGAACTTTAAAATTAGACATAATGTGGAGGATGGGAGCGTGCAACTAGCGGATCATTATCAACA |
| | GAATACGCCTATTGGTGATGGCCCCGTGCTGCTTCCCGATAATCATTATCTAAGCACACAAAGCGTGTT |
| | GAGCAAAGATCCAAATGAGAAACGGGATCATATGGTGTTGCTCGAATTTGTCACAGCCGCGGGAATTAC |
| | ACATGGAATGGATGAGTTGTATAAA |
| GFP25 | ATGAGCAAAGGTGAAGAACTGTTTACCGGCGTTGTGCCGATTCTGGTGGAACTTGATGGTGATGTGAAT |
| | GGGCATAAATTTAGCGTCAGGGGCGAAGGAGAGGGTGATGCGACGAACGGAAAACTGACGCTGAAATT |
| | TATTTGTACCACCGGTAAACTTCCGGTCCCATGGCCCACCCTCGTGACCACACTAACCTATGGAGTCCAA |
| | TGTTTTAGCCGCTATCCTGATCATATGAAACGCCATGATTTTTTTAAAAGCGCGATGCCTGAGGGCTAT |
| | GTGCAAGAGCGTACCATTAGCTTTAAAGATGATGGAACCTATAAAACCCGAGCGGAAGTCAAATTTGAG |
| | GGCGATACCCTAGTGAACAGAATTGAGCTGAAAGGAATTGATTTTAAAGAGGATGGCAATATTTTAGGG |
| | CATAAACTCGAGTATAATTTCAACAGCCATAATGTGTATATTACCGCCGATAAACAAAAAAATGGAATC |
| | AAAGCGAACTTTAAAATTAGGCACAACGTGGAAGATGGAAGCGTGCAACTTGCGGATCATTATCAGCAG |
| | AATACACCGATTGGTGATGGCCCAGTGCTCCTCCCAGATAATCATTATCTTAGCACCCAGAGCGTCCTGA |
| | GCAAAGATCCAAATGAGAAAAGGGATCATATGGTGCTGCTTGAGTTTGTTACCGCCGCGGGGATTACGC |
| | ACGGAATGGATGAGTTATATAAA |
| GFP50 | ATGAGCAAAGGTGAAGAACTGTTTACCGGCGTTGTGCCGATTCTGGTGGAACTGGATGGTGATGTGAAT |
| | GGCCATAAATTTAGCGTTCGTGGCGAAGGAGAAGGAGATGCGACCAACGGTAAACTGACCCTGAAATTT |
| | ATTTGCACCACCGGTAAACTGCCGGTTCCGTGGCCCACCCTGGTGACCACCCTGACCTATGGCGTTCAGT |
| | GTTTTAGCCGCTATCCGGATCATATGAAACGCCATGATTTCTTTAAAAGCGCGATGCCGGAAGGCTATG |
| | TGCAGGAACGTACCATTAGCTTCAAAGATGATGGCACCTATAAAACCCGTGCGGAGGTGAAATTTGAAG |
| | GCGATACCCTCGTGAACCGCATTGAACTGAAAGGTATTGATTTTAAAGAAGATGGCAATATTCTGGGTC |
| | ATAAACTGGAATATAATTTTAACAGCCATAATGTGTATATTACCGCCGATAAACAGAAAAATGGCATCA |
| | AAGCGAATTTTAAAATCCGTCACAACGTGGAGGATGGTAGCGTGCAGTTAGCGGATCATTATCAGCAGA |
| | ATACCCCGATTGGTGATGGCCCGGTGCTGCTGCCGGATAATCATTATCTGAGCACCCAAAGCGTTCTGA |
| | GCAAAGATCCGAATGAAAAACGTGATCATATGGTGCTGCTGGAATTTGTTACCGCCGCGGGCATTACCC |
| | ACGGTATGGATGAACTGTATAAA |
| GFP75 | ATGAGCAAAGGTGAAGAACTGTTTACCGGCGTTGTGCCGATTCTGGTGGAACTGGATGGTGATGTTAAC |
| | GGCCACAAATTCAGCGTTCGTGGCGAAGGCGAAGGTGACGCTACCAACGGTAAACTGACCCTGAAATTT |
| | ATCTGCACCACCGGTAAACTGCCGGTTCCGTGGCCGACCCTGGTGACCACCCTGACCTATGGCGTTCAG |
| | TGCTTCAGCCGCTACCCGGACCATATGAAACGCCATGACTTCTTCAAAAGCGCGATGCCGGAAGGCTAT |
| | GTGCAGGAACGTACCATTAGCTTCAAAGATGACGGCACCTATAAAACCCGTGCGGAAGTTAAATTCGAA |
| | GGCGATACCCTGGTTAACCGCATCGAACTGAAAGGTATCGATTTTAAAGAAGACGGCAACATCCTGGGT |
| | CATAAACTGGAATATAACTTCAACAGCCACAATGTTTATATTACCGCCGATAAACAGAAAAACGGCATCA |
| | AAGCGAACTTTAAAATCCGTCACAACGTAGAAGATGGTTCTGTGCAGCTGGCGGATCATTATCAGCAGA |
| | ACACCCCGATCGGTGATGGCCCGGTACTGCTGCCGGATAATCATTACCTGAGCACCCAGTCCGTTCTGT |
| | CTAAAGATCCGAATGAAAAACGTGACCATATGGTGCTGCTGGAATTTGTTACCGCTGCTGGCATCACCC |
| | ACGGTATGGACGAACTGTACAAA |
| GFP90 | ATGAGCAAAGGTGAAGAACTGTTTACCGGCGTTGTGCCGATTCTGGTGGAACTGGACGGTGATGTTAAC |
| | GGCCACAAATTCAGCGTTCGTGGCGAAGGCGAAGGTGACGCTACCAACGGTAAACTGACCCTGAAATTC |
| | ATCTGCACCACCGGTAAACTGCCGGTTCCGTGGCCGACCCTGGTAACCACCCTGACCTATGGCGTTCAG |
| | TGCTTCAGCCGCTACCCGGACCACATGAAACGCCACGACTTCTTCAAATCTGCTATGCCGGAAGGCTATG |
| | TACAGGAACGTACCATCAGCTTCAAAGACGACGGCACCTACAAAACCCGTGCGGAAGTTAAATTCGAAG |
| | GCGACACCCTGGTTAACCGCATCGAACTGAAAGGTATCGATTTCAAAGAAGACGGCAACATCCTGGGTC |
| | ACAAACTGGAATACAACTTCAACTCTCACAACGTTTACATTACCGCCGACAAACAGAAAAACGGCATCAA |
| | AGCTAACTTCAAAATCCGTCACAACGTAGAAGACGGTTCTGTACAGCTGGCGGACCATTACCAGCAGAA |
| | CACCCCGATCGGTGATGGCCCGGTACTGCTGCCGGACAACCACTACCTGAGCACCCAGTCCGTTCTGTCT |
| | AAAGATCCGAACGAAAAACGTGACCACATGGTTCTGCTGGAATTTGTTACCGCTGCTGGCATCACCCAC |
| | GGTATGGACGAACTGTACAAA |

Supplementary Table S5: **Sequences of mCherry2 genes with different codon usage**

| CDS Name | Nucleotide Sequence |
|---|---|
| MCH10 | ATGGTGAGCAAGGGCGAGGAGGATAACCTGGCCATCATCAAGGAGTTTATGCGGTTTAAGGTGCATATG<br>GAGGGGAGTGTGAATGGACATGAGTTCGAGATCGAGGGGGAGGGAGAGGGCAGACCCTATGAGGGAAC<br>ACAAACGGCCAAGTTGAAGGTGACGAAGGGTGGACCCTTGCCCTTTGCCTGGGATATTCTCAGTCCTCA<br>ATTTATGTATGGAGCAAGGCCTATGTGAAGCATCCCGCCGACATACCCGATTACTTGAAGTTGAGTTT<br>TCCCGAGGGGTTTAATTGGGAGAGGGTGATGAATTTTGAGGATGGCGGGGTGGTGACCGTGACCCAGG<br>ATTCAAGTCTTCAAGATGGCGAGTTTATATATAAGGTGAAGTTACGAGGGACAAATTTTCCCAGCGATG<br>GACCCGTAATGCAATGTCGAACCATGGGGTGGGAGGCCTCGACAGAGCGGATGTACCCCGAGGATGGAG<br>CCTTGAAGGGAGAGATTAAGCAAAGGCTTAAGTTAAAGGATGGGGGACATTATGATGCCGAGGTCAAGA<br>CGACATATAAGGCCAAGAAGCCCGTGCAACTACCCGGAGCCTATAATGTCGATATAAAGTTGGATATAC<br>TTTCGCATAATGAGGACTATACGATTGTGGAACAATATGAGCGGGCCGAGGGGCGGCATAGCACGGGAG<br>GGATGGATGAGCTTTACAAG |
| MCH25 | ATGGTGAGCAAGGGCGAGGAGGATAACCTGGCCATCATCAAGGAGTTCATGAGATTTAAGGTGCATATG<br>GAGGGCTCAGTGAATGGCCACGAGTTTGAGATTGAGGGCGAGGGCGAGGGGCGCCCCTATGAGGGCAC<br>ACAAACAGCCAAGTTGAAGGTGACCAAGGGAGGGCCCTTACCCTTCGCCTGGGATATATTGTCACCTCA<br>ATTTATGTATGGGTCGAAGGCCTATGTGAAGCACCCCGCCGATATCCCCGACTACTTGAAGCTGTCATT<br>TCCCGAGGGGTTTAATTGGGAGCGGGTGATGAATTTCGAGGACGGCGGGGTGGTGACCGTGACCCAAG<br>ATAGTAGTCTGCAAGATGGCGAGTTTATATATAAGGTGAAGTTGAGAGGAACCAATTTCCCCTCGGACG<br>GCCCCGTGATGCAATGTAGGACAATGGGCTGGGAGGCCAGCACTGAGCGGATGTATCCCGAGGACGGAG<br>CCTTAAAGGGAGAGATCAAGCAGAGGCTGAAGCTTAAGGACGGAGGGCATTACGACGCCGAGGTCAAG<br>ACAACCTACAAGGCCAAGAAGCCCGTGCAATTGCCCGGCGCCTATAATGTCGATATCAAGTTGGATATT<br>CTTTCCCACAATGAGGACTATACCATCGTGGAACAGTACGAGCGGGCCGAGGGAAGGCATAGTACAGGC<br>GGCATGGATGAGCTCTATAAG |
| MCH50 | ATGGTGAGCAAGGGCGAGGAGGATAACCTGGCCATCATCAAGGAGTTCATGAGATTCAAGGTGCACATG<br>GAGGGCTCCGTGAACGGCCATGAGTTCGAGATCGAGGGCGAGGGCGAGGGCCGCCCCTATGAGGGCAC<br>ACAGACCGCCAAGCTTAAGGTGACAAAGGGTGGCCCCCTGCCCTTCGCCTGGGACATACTGAGTCCTCA<br>GTTCATGTACGGCTCCAAGGCCTACGTGAAGCACCCCGCCGACATCCCCGACTACTTGAAGCTGTCCTTC<br>CCCGAGGGCTTTAATTGGGAGCGCGTGATGAACTTCGAGGACGGCGGCGTGGTGACCGTGACCCAGGAC<br>TCCTCCCTGCAGGACGGCGAGTTTATCTACAAGGTGAAGCTGCGCGGCACCAACTTCCCCTCCGATGGA<br>CCCGTAATGCAGTGTAGAACAATGGGATGGGAGGCCTCCACAGAGCGGATGTACCCCGAGGACGGCGCC<br>CTGAAGGGGGAGATCAAGCAGAGGTTGAAGCTTAAGGACGGGGGCCACTACGACGCTGAGGTCAAGAC<br>CACCTACAAGGCCAAGAAGCCCGTGCAACTGCCCGGGGCCTACAACGTCGACATCAAGTTGGACATCCT<br>TAGTCACAACGAGGACTACACCATCGTGGAGCAATATGAGCGCGCCGAGGGAAGGCATTCCACCGGCGG<br>CATGGACGAGTTATACAAGTAATAA |
| MCH75 | ATGGTGAGCAAGGGCGAGGAGGATAACCTGGCCATCATCAAGGAGTTCATGCGCTTCAAGGTGCACATG<br>GAGGGCTCCGTGAACGGCCACGAGTTCGAAATCGAAGGCGAGGGCGAGGGCCGCCCCTACGAGGGCACC<br>CAGACCGCCAAGCTGAAGGTGACCAAGGGTGGCCCCCTGCCCTTCGCTTGGGACATCCTGTCCCCTCAG<br>TTCATGTACGGCTCCAAGGCCTACGTGAAGCACCCCGCCGACATCCCCGACTACTTGAAGCTGTCCTTCC<br>CCGAGGGCTTCAATTGGGAGCGCGTAATGAACTTCGAAGACGGCGGCGTGGTGACCGTTACCCAGGACT<br>CCTCCCTGCAGGACGGCGAGTTCATCTACAAGGTGAAGCTGCGCGGCACCAACTTCCCCTCCGACGGCC<br>CCGTAATGCAGTGCCGTACCATGGGCTGGGAAGCTTCCACTGAACGGATGTACCCCGAGGACGGCGCCC<br>TGAAGGGCGAGATCAAGCAGAGGCTGAAGCTGAAGGACGGCGGCCACTACGACGCTGAAGTCAAGACCA<br>CCTACAAGGCCAAGAAGCCCGTTCAGCTGCCGGGCGCCTACAACGTCGACATCAAGCTGGACATCCTTT<br>CCCACAACGAGGACTACACCATCGTGGAACAGTACGAACGCGCTGAAGGCCGCCACTCCACCGGCGGCA<br>TGGACGAACTGTACAAG |
| MCH90 | ATGGTGAGCAAGGGCGAGGAGGATAACCTGGCCATCATCAAGGAATTCATGCGTTTCAAAGTCCACATG<br>GAGGGTTCTGTAAACGGCCACGAATTCGAAATCGAAGGTGAAGGTGAAGGTCGTCCGTACGAAGGCACT<br>CAGACTGCTAAGCTGAAAGTTACCAAAGGCGGTCCGCTGCCGTTCGCTTGGGACATCCTGTCCCCGCAG<br>TTCATGTACGGTTCCAAAGCTTACGTAAAGCACCCGGCCGACATACCGGACTACCTAAAGTTGTCTTTCC<br>CGGAAGGTTTCAACTGGGAACGCGTTATGAACTTCGAAGACGGCGGCGTTGTCACTGTAACTCAGGACT<br>CTTCTTTACAGGACGGTGAATTCATCTACAAGGTAAAGCTGCGCGGTACCAACTTCCCTTCTGACGGTCC<br>GGTAATGCAGTGCCGTACTATGGGCTGGGAAGCTTCTACCGAAAGGATGTACCCGGAAGACGGTGCTCT<br>GAAAGGTGAAATCAAGCAGCGTTTAAAGCTGAAGGACGGCGGCCACTACGACGCTGAAGTTAAGACCAC<br>TTACAAAGCTAAGAAACCGGTTCAGCTGCCGGGCGCTTACAACGTTGACATCAAGCTGGACATCCTGTC<br>TCACAACGAAGACTACACTATCGTTGAACAGTACGAACGTGCTGAAGGTCGTCACAGTACTGGCGGCAT<br>GGACGAATTGTATAAG |

Supplementary Table S6: **Sequences of sfGFP genes with one target codon deoptimized**

| CDS Name | Nucleotide Sequence |
|---|---|
| GFP GGG | ATGAGCAAAGGTGAAGAACTGTTTACCGGCGTTGTGCCGATTCTGGTGGAACTGGATGGGGATGTGAAT<br>GGGCACAAATTTAGCGTTCGTGGGGAAGGGGAAGGGGATGCGACCAACGGGAAACTGACCCTGAAATTT<br>ATTTGCACCACCGGGAAACTGCCGGTTCCGTGGCCCACCCTGGTGACCACCCTGACCTATGGGGTTCAG<br>TGTTTTAGCCGCTATCCGGATCATATGAAACGCCATGATTTCTTTAAAAGCGCGATGCCGGAAGGGTAT<br>GTGCAGGAACGTACCATTAGCTTCAAAGATGATGGGACCTATAAAACCCGTGCGGAGGTGAAATTTGAA<br>GGGGATACCCTCGTGAACCGCATTGAACTGAAAGGGATTGATTTTAAAGAAGATGGGAATATTCTGGGG<br>CATAAACTGGAATATAATTTTAACAGCCATAATGTGTATATTACCGCCGATAAACAGAAAAATGGGATC<br>AAAGCGAATTTTAAAATCCGTCACAACGTGGAGGATGGGAGCGTGCAGTTAGCGGATCATTATCAGCAG<br>AACACCCCGATTGGGGATGGGCCGGTGCTGCTGCCGGATAATCATTATCTGAGCACCCAATCCGTTCTG<br>AGCAAAGATCCGAATGAAAAACGTGATCATATGGTGCTGCTGGAATTTGTTACCGCCGCGGGGATTACC<br>CACGGGATGGATGAACTGTATAAA |
| GFP GGA | ATGAGCAAAGGTGAAGAACTGTTTACCGGCGTTGTGCCGATTCTGGTGGAACTGGATGGAGATGTGAAT<br>GGACACAAATTTAGCGTTCGTGGAGAAGGAGAAGGAGATGCGACCAACGGAAAACTGACCCTGAAATTT<br>ATTTGCACCACCGGAAAACTGCCGGTTCCGTGGCCCACCCTGGTGACCACCCTGACCTATGGAGTTCAG<br>TGTTTTAGCCGCTATCCGGATCATATGAAACGCCATGATTTCTTTAAAAGCGCGATGCCGGAAGGATAT<br>GTGCAGGAACGTACCATTAGCTTCAAAGATGATGGAACCTATAAAACCCGTGCGGAGGTGAAATTTGAA<br>GGAGATACCCTCGTGAACCGCATTGAACTGAAAGGAATTGATTTTAAAGAAGATGGAAATATTCTGGGA<br>CATAAACTGGAATATAATTTTAACAGCCATAATGTGTATATTACCGCCGATAAACAGAAAAATGGAATC<br>AAAGCGAATTTTAAAATCCGTCACAACGTGGAGGATGGAAGCGTGCAGTTAGCGGATCATTATCAGCAG<br>AACACCCCGATTGGAGATGGACCGGTGCTGCTGCCGGATAATCATTATCTGAGCACCCAATCCGTTCTG<br>AGCAAAGATCCGAATGAAAAACGTGATCATATGGTGCTGCTGGAATTTGTTACCGCCGCGGGAATTACC<br>CACGGAATGGATGAACTGTATAAA |
| GFP GAG | ATGAGCAAAGGTGAAGAACTGTTTACCGGCGTTGTGCCGATTCTGGTGGAGCTGGATGGTGATGTGAAT<br>GGCCATAAATTTAGCGTTCGTGGCGAGGGAGAGGGAGATGCGACCAACGGTAAACTGACCCTGAAATTT<br>ATTTGCACCACCGGTAAACTGCCGGTTCCGTGGCCCACCCTGGTGACCACCCTGACCTATGGCGTTCAGT<br>GTTTTAGCCGCTATCCGGATCATATGAAACGCCATGATTTCTTTAAAAGCGCGATGCCGGAGGGCTATG<br>TGCAGGAGCGTACCATTAGCTTCAAAGATGATGGCACCTATAAAACCCGTGCGGAGGTGAAATTTGAGG<br>GCGATACCCTCGTGAACCGCATTGAGCTGAAAGGTATTGATTTTAAAGAGGATGGCAATATTCTGGGTC<br>ATAAACTGGAGTATAATTTTAACAGCCATAATGTGTATATTACCGCCGATAAACAGAAAAATGGCATCA<br>AAGCGAATTTTAAAATCCGTCACAACGTGGAGGATGGTAGCGTGCAGTTAGCGGATCATTATCAGCAGA<br>ACACCCCGATTGGTGATGGCCCGGTGCTGCTGCCGGATAATCATTATCTGAGCACCCAATCCGTTCTGA<br>GCAAAGATCCGAATGAGAAACGTGATCATATGGTGCTGCTGGAGTTTGTTACCGCCGCGGGCATTACCC<br>ACGGTATGGATGAGCTGTATAAA |
| GFP CTA | ATGAGCAAAGGTGAAGAACTGTTTACCGGCGTTGTGCCGATTCTGGTGGAACTAGATGGTGATGTGAAT<br>GGCCACAAATTTAGCGTTCGTGGCGAAGGAGAAGGAGATGCGACCAACGGTAAACTAACCCTAAAATTT<br>ATTTGCACCACCGGTAAACTACCGGTTCCGTGGCCCACCCTAGTGACCACCCTAACCTATGGCGTTCAGT<br>GTTTTAGCCGCTATCCGGATCATATGAAACGCCATGATTTCTTTAAAAGCGCGATGCCGGAAGGCTATG<br>TGCAGGAACGTACCATTAGCTTCAAAGATGATGGCACCTATAAAACCCGTGCGGAGGTGAAATTTGAAG<br>GCGATACCCTAGTGAACCGCATTGAACTAAAAGGTATTGATTTTAAAGAAGATGGCAATATTCTAGGTC<br>ATAAACTAGAATATAATTTTAACAGCCATAATGTGTATATTACCGCCGATAAACAGAAAAATGGCATCA<br>AAGCGAATTTTAAAATCCGTCACAACGTGGAGGATGGTAGCGTGCAGCTAGCGGATCATTATCAGCAGA<br>ACACCCCGATTGGTGATGGCCCGGTGCTACTACCGGATAATCATTATCTAAGCACCCAATCCGTTCTAAG<br>CAAAGATCCGAATGAAAAACGTGATCATATGGTGCTACTAGAATTTGTTACCGCCGCGGGCATTACCCA<br>CGGTATGGATGAACTATATAAA |
